# Heterologous expression of insect IRs in transgenic *Drosophila melanogaster*

**DOI:** 10.1101/2023.09.12.557369

**Authors:** Alberto Maria Cattaneo, Cristina M. Crava, William B Walker

## Abstract

Among insect chemosensory receptors, the broadly conserved Ionotropic Receptors (IRs) remain some of the least investigated. Several studies have documented IR activation by recording insect’s neuronal activity *in situ*, some demonstrated their activation when expressed in oocytes from *Xenopus*, others made use of the *Drosophila* “ionotropic receptor decoders” to functionally mis-express IRs from the same species or from the closely related *D. sechellia*. Here we demonstrated that both substituting tuning IRs of *D. melanogaster* and expressing heterologous IRs from other insects alongside the *Drosophila* native ones result in functional heteromeric complexes. By these methods, we functionally characterized the IR41a1 subunit of the codling moth *Cydia pomonella*, which demonstrated binding to polyamines with different pharmacological characteristics and the IR75d subunit of the spotted wing drosophila *Drosophila suzukii*, which binds hexanoic acid. Then we expressed the *D. suzukii* acid sensor IR64a into the *D. melanogaster* “ionotropic receptor decoder” neuron, which resulted in the inhibition to the main activators of other neurons housed in the same sensillum of *D. melanogaster*, as an evidence of its possible functional expression, but it did not show response to acids. *In situ* hybridization on the antennae of *D. suzukii* unveiled a wide expression of this subunit in neurons proximal to the sacculus. Structural analysis did not explain absence of IR64a binding to acids, but it identified key amino acid features that may justify possible hexanoic acid binding for IR75d. While our findings add to the derophanization efforts conducted on the chemosensors of the aforementioned pests, they demonstrated potential for the use of neurons of transgenic *Drosophila* as a tool to functionally characterize IRs from different insect species.

## Introduction

Ionotropic Receptors (IRs) are transmembrane chemoreceptors found in the peripheral sensory neurons of animals belonging to the superphylum Protostomia, which includes arthropods, nematodes, mollusks, annelids and other invertebrates (Wicher and Miazzi 2021, Eyun et al., 2017, Croset et al., 2010). These receptors share an evolutionary relationship to the Ionotropic Glutamate Receptors (iGluRs), which are an ancient class of ligand-gated ion channels involved in neuronal communication across the animal kingdom (Gereau & Swanson, 2008). iGluRs are also present in a small number of prokaryotes (Chen et al., 1999; Chiu et al., 1999) and plants (Lam et al., 1998).

IRs were first discovered in *Drosophila melanogaster* (Benton et al., 2009), and subsequent research on IRs has primarily focused on this species and a few mosquito species. IRs function as ligand-gated ion channels, allowing the influx of cations into the cytoplasm and activation of the olfactory sensory neuron (OSN) (Abuin et al., 2019). Each IR-gated ion channel consists of individual odor-specific tuning subunits and one or two broadly expressed co-receptors (Ir25a, Ir8a, and Ir76b) (Abuin et al., 2011, Abuin et al., 2019, Vulpe and Menuz, 2021). In *D. melanogaster*, the functional characterization of the tuning IR subunits led to the identification of agonists activating nine subunits (Ni, 2021). Functional characterization in *D. melanogaster* has benefitted from the genetic toolbox available for this species, which enables a variety of approaches. These include recording ligand-induced neuronal activity *in situ* in knocked out lines (Ai et al., 2010; Grosjean et al., 2011; Min et al., 2013; Gorter el et., 2016), functional imaging experiments (Silbering et al., 2011; Hussain et al., 2016; Ai et al., 2013; Min et al., 2013) or mis-expression of heterologous IRs in different OSNs (Benton et al, 2009; Abuin et al., 2011; Ai et al, 2013; Abuin et al., 2019) or in the “ionotropic receptor decoder”, which is an OSN lacking the endogenous tuning receptor subunit but expressing the IR8a coreceptor (Prieto-Godino 2016; 2017; 2021, Abuin et al; 2011, Grosjean 2011). In contrast, similar genetic tools are not readably available for non-model species, slowing down then comprehension of the role of IRs outside *Drosophila*. So far, the methods used include knock down and *in situ* recording of neuronal activity (Xu et al, 2014) or the heterologous expression in *Xenopus laevis* oocytes (Xu et al, 2014; Ray et al., 2023; Pitts et al, 2017; Shan et al 2019; Hou et al 2022). This is a commonly used heterologous expression system for the electrophysiological characterization of iGluRs and it has been the only method used for functional characterization of IRs from lineages outside Diptera species, specifically the lepidopteran *Agrotis segetum* (Hou et al., 2022) and the wasp *Microplitis mediator* (Shan et al; 2019). However, it relies on a single cell system extremely different from the native OSN environment, which imposes significant limitations on our understanding of the activation and pharmacology of the expressed receptor (Hou et al., 2020).

An interesting *in vivo* alternative is the “ionotropic receptor decoder”, which closely resembles the *Drosophila* “empty-neuron” system. The latter has been extensively used for functional characterization of another class of insect olfactory receptors, the odorant receptors (ORs), in insect species within and beyond dipterans (Gonzalez et al., 2016). In contrast, the “ionotropic receptor decoder” has been successfully used to deorphanize IR tuning subunits only from *D. melanogaster* and from a closely related species from the melanogaster subgroup, *Drosophila sechellia* (Prieto-Godino 2016; 2017; 2021, Abuin et al; 2011, Grosjean 2011). Additionally, the “ionotropic receptor decoder” is restricted to IR tuning subunits that function with the IR8a co-receptor, since it relies on an OSN housed in ac4 sensilla and not expressing the native IR84 tuning subunit. In contrast, heterologous expressions in OSNs co-expressing IR25a and IR76b have not been assessed.

In this study, our aim is to explore alternative methods for heterologous expression of IRs, with the goal of providing new tools for comprehensive functional characterization of insect IRs beyond *Drosophila*. Specifically, we focused on two horticultural pests: the lepidopteran codling moth, *Cydia pomonella* and one drosophilid species outside the melanogaster species subgroup, the spotted winged drosophila, *Drosophila suzukii*. In both species, we selected IRs orthologous to *D. melanogaster* DmelIR41a, DmellIR75d, and DmelIR64a, whose agonists are already known (Hussain et al., 2016; Silbering et al., 2011; Ai et al., 2010; Ai et al., 2013). To achieve this goal, we explored the use of two systems: an *in vivo* expression system using several OSNs housed in the ac4 sensillum of *D. melanogaster* as target and an *ex vivo* heterologous system utilizing Human Embryonic Kidney (HEK293T) cells. The ac4 sensillum houses three OSNs, each expressing one of three tuning subunits: DmelIR84a, DmelIR75d, and DmelIR76a (Silbering et al, 2011) (Figure 1). The responses of both DmelIR75d and DmelIR76a are dependent on co-receptor DmelIR25a, although the latter also requires the expression of the DmelIR76b co-receptor subunit for accurate detection of polyamines (Siblering et, 2011, Abuin et al, 2011). On the contrary, DmelIR84a subunit forms a heterotetramer with DmelIR8a co-receptor (Abuin et al., 2011; Grosjean et al., 2011; Silbering et al., 2011). We substituted the tuning receptors with *D. suzukii* IRs or expressed the heterologous *C. pomonella* tuning receptors alongside the native receptors, demonstrating that both approaches result in a functional heteromeric complex. In contrast, our attempts to express functional IRs in HEK293T cells were unfruitful. Our results provide the first successful examples of heterologous *in vivo* expression in *Drosophila* of IRs from distant species either by use of novel “ionotropic receptor decoders” or by expressing a heterologous IR alongside a native one. This breakthrough opens up new possibilities for the application of these powerful tools in the deorphanization of ionotropic receptors of non-model insect species.

**Figure 1.**
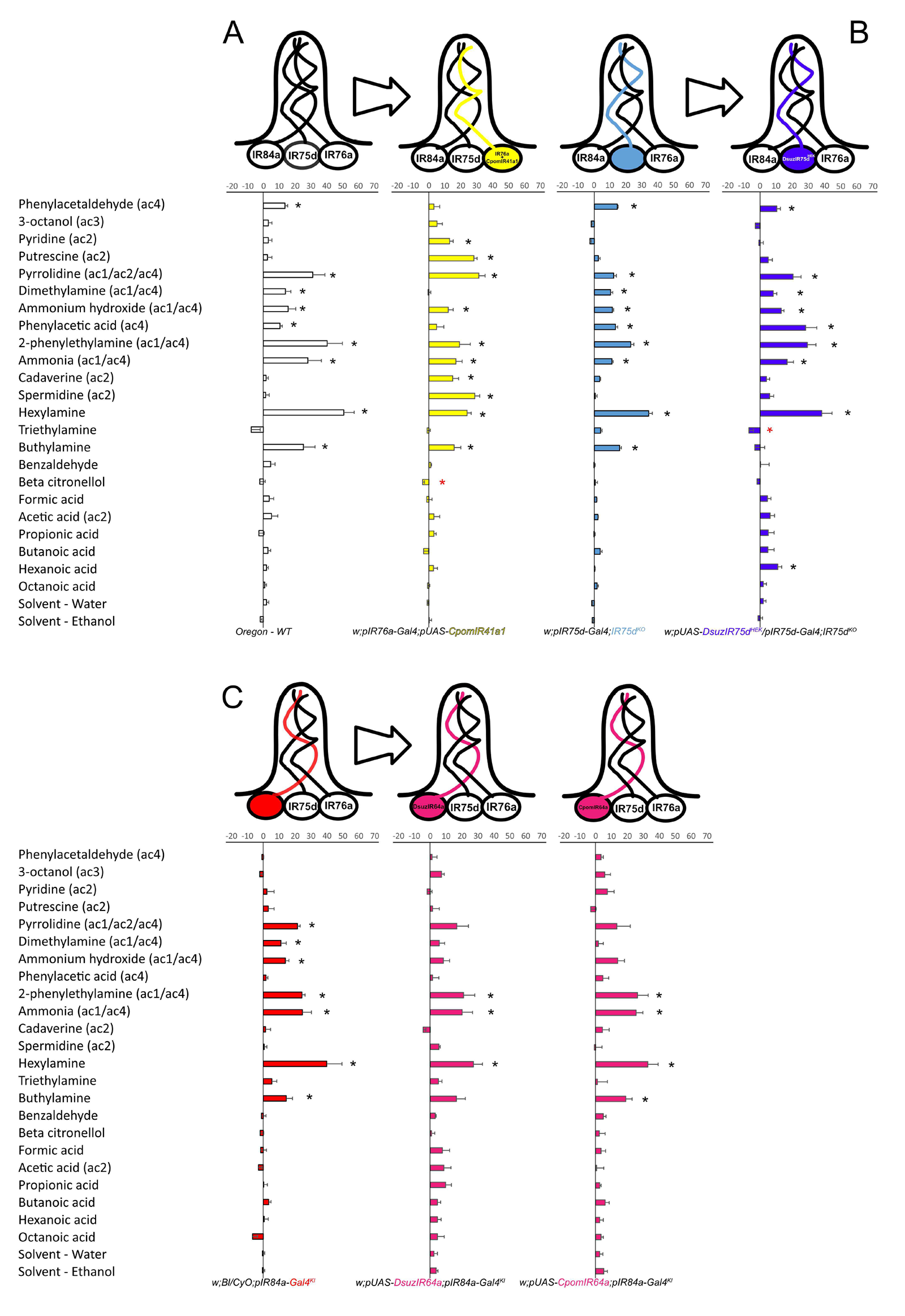
Heterologous expression of *C. pomonella* and *D. suzukii* IRs in ac4-neurons of *D. melanogaster*. (**A**) – Odor response profiles from ac4-sensilla of *D. melanogaster* comparing Oregon-C wild type (white) with the transgenic fly line expressing *CpomIR41a1* of *C. pomonella* in IR76a neurons [*w;pIR76a-Gal4;pUAS-CpomIR41a1* – yellow]. (**B**) – Odor response profiles from ac4-sensilla of *D. melanogaster* comparing *IR75d^KO^*flies [*w;pIR75d-Gal4;IR75d^KO^* – light blue] with the transgenic fly line expressing *DsuzIR75d^HEK^* of *D. suzukii* [*w;pIR75d-Gal4/pUAS-DsuzIR75d^HEK^;IR75d^KO^* – blue]. (**C**) – Odor response profiles from ac4-sensilla of *D. melanogaster* comparing *IR84a^KO^* flies [*w;Bl/CyO;pIR84a-Gal4^KI^*-red] with the transgenic fly lines expressing either *DsuzIR64a* of *D. suzukii* [*w;pUAS-DsuzIR64a;pIR84a-Gal4*] or the truncated *CpomIR64a* isoform of *C. pomonella* [*w;pUAS-CpomIR64a;pIR84a-Gal4*] (magenta). The compound library tested is reported in Table 1. Asterisks indicate compounds enhancing significant increment (black) or decrement (red) in spike frequency (Δspikes/0.5 s) when compared with their respective solvent, water or ethanol, in a paired T-test (p < 0.05; two tails, N = 5-9 depending on the experiment; Supplementary data file 1). Values in parenthesis (ac1-4) indicate whether the compound is active on a specific type of coeloconic (ac) sensilla, based on Silbering et al., (2011) and on the database of odorant responses (http://neuro.uni-konstanz.de/DoOR/content/DoOR.php; Munch and Galizia (2016), Galizia et al., (2010)) (Table 1).

## Results

### Functional characterization of C. pomonella IR41a1 heterologously expressed in D. melanogaster olfactory sensory neuron alongside a native IR

We heterologously expressed *C. pomonella* IR41a1 (CpomIR41a1) in the ac4 sensillum of *D. melanogaster* using a pIR76a-Gal4 driver line (Silbering et al., 2011) (Figure 1). Consequently, CpomIR41a1 was co-expressed in the same OSN along with the native IR76a tuning subunit (DmelIR76a), and the broadly-expressed co-receptors DmelIR25a and DmelIR76b. DmelIR76a’s best-known agonists are pyrrolidine and phenylethylamine (Abuin et al., 2011; Silbering et al., 2011). When we use single sensillum recording to quantify the spiking response of the mutant ac4 sensilla expressing CpomIR41a, testing compounds from Table 1 unveiled response to two ligands quantitatively similar to the control ac4 sensilla (pyrrolidine, *p* < 0,001 and 2-phenylethylamine, *p* = 0.017). This suggested that expression of a second IRs in the same OSN was not interfering with the activity of the native one. However, CpomIR41a1 conferred robust responses to pyridine (*p* = 0.005), putrescine (*p* < 0.001), cadaverine (*p* = 0.003) and spermidine (*p* < 0.001) compared to control neurons that do not express this tuning IR (Figure 1, Supplementary data file 1), indicating that these four ligands are agonist of CpomIR41a1. Additionally, a slight inhibition of spiking activity in ac4 sensilla was observed with beta citronellol (-3,85 ± 0,97 spikes/0.5s; p = 0.019), indicating a potential role of this volatile compound as antagonist of CpomIR41a1. Interestingly, expression of CpomIR41a1 abolished dimethylamine-evoked response (p = 0.893) as well as responses to phenylacetaldehyde (p = 0.116) and phenylacetic acid (p = 0.167), which are stimulus known to activate IR84a-expressing OSNs. This suggests that the expression of CpomIR41a1 may interfere with the proper response of other OSNs housed in the same sensillum.

**Table 1.**
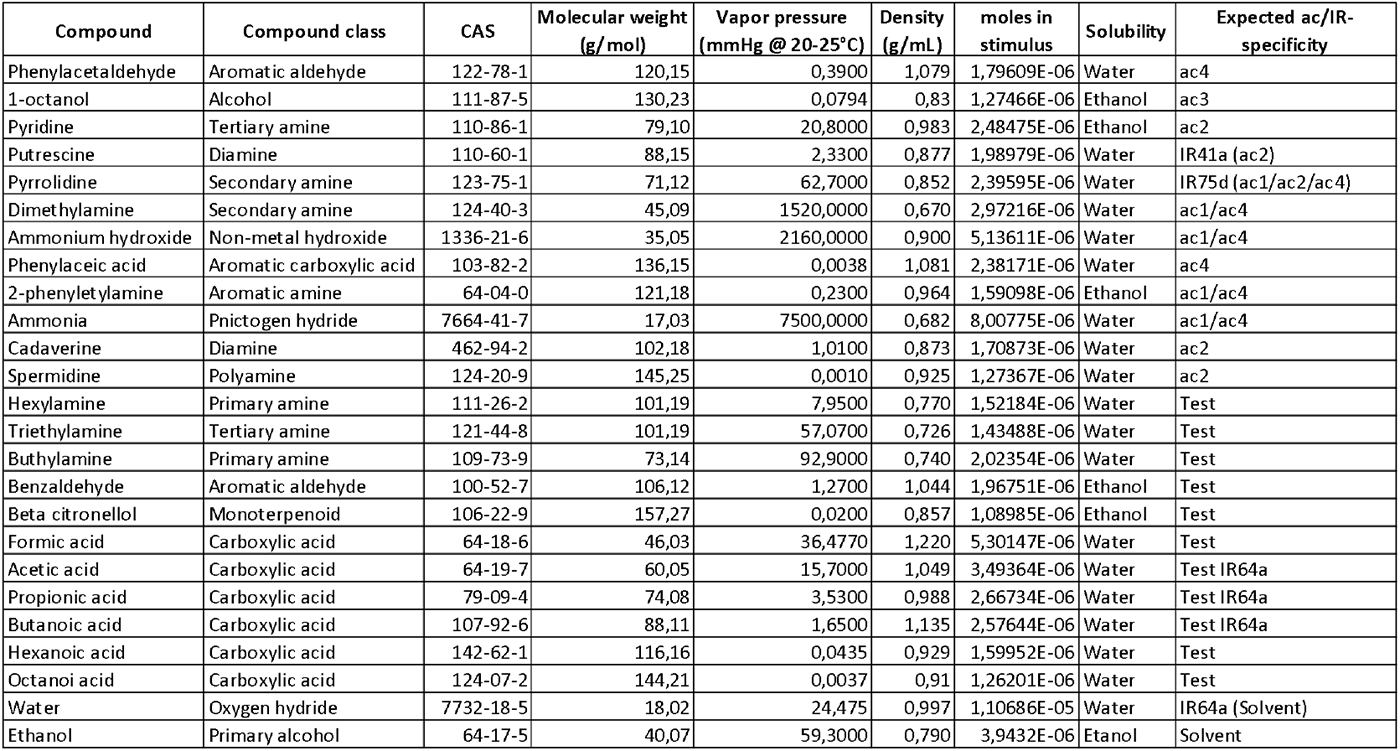
Panel of ligands tested on transgenic *D. melanogaster*. Chemical and physical properties of the tested synthetic ligands were obtained consulting the database of odorant responses (http://neuro. uni-konst anz. de/ DoOR/ conte nt/ DoOR. php; Munch and Galizia (2016), Galizia et al., (2010)) and the good scents company (http://www.thegoodscentscompany.com/search2.html). Moles to the tested doses were based on using 20 μL of 1.0% dilutions or 30 μL of 10 μg/μL for phenylacetic acid. The expected specificity for ac-sensilla or IRs was based on reported data from Silbering et al., (2011) and from the database of odorant responses.

| Compound | Compound class | CAS | Molecular weight (g/mol) | Vapor pressure (mmHg @ 20-25°C) | Density (g/ml) | moles in stimulus | Solubility | Expected ac/IR specificity |
| --- | --- | --- | --- | --- | --- | --- | --- | --- |
| Phenylacetaldehyde | Aromatic aldehyde | 122-78-1 | 120,15 | 0,3900 | 1,079 | 1,79 609E-06 | Water | ac4 |
| 1-octanol | Alcohol | 111-87-5 | 130,23 | 0,0794 | 0,83 | 1,27466E-06 | Ethanol | ac3 |
| Pyridine | Tertiary amine | 110-86-1 | 79,10 | 20,8000 | 0,983 | 2,48475E-06 | Ethanol | ac2 |
| Putrescine | Diamine | 110-60-1 | 88,15 | 2,3300 | 0,877 | 1,98979E-06 | Water | IR41a (ac2) |
| Pyrrolidine | Secondary amine | 123-75-1 | 71,12 | 62,7000 | 0,852 | 2,39 595E-06 | Water | IR75d (ac1/ac2/ac4) |
| Dimethylamine | Secondary amine | 124-40-3 | 45,09 | 1520,0000 | 0,670 | 2,97216E-06 | Water | ac1/ac4 |
| Ammonium hydroxide | Non-metal hydroxide | 1336-21-6 | 35,05 | 2160,0000 | 0,900 | 5,13611E-06 | Water | ac1/ac4 |
| Phenylacetic acid | Aromatic carboxylic acid | 103-82-2 | 136,15 | 0,0038 | 1,081 | 2,38 171E-06 | Water | ac4 |
| 2-phenylethylamine | Aromatic amine | 64-04-0 | 121,18 | 0,2300 | 0,964 | 1,59098E-06 | Ethanol | ac1/ac4 |
| Ammonia | pnictogen hydride | 7664-41-7 | 17,03 | 7500,0000 | 0,682 | 8,00775E-06 | Water | ac1/ac4 |
| Cadaverine | Diamine | 462-94-2 | 102,18 | 1,0100 | 0,873 | 1,70873E-06 | Water | ac2 |
| Spermidine | Polyamine | 124-20-9 | 145,25 | 0,0010 | 0,925 | 1,27367E-06 | Water | ac2 |
| Hexylamine | Primary amine | 111-26-2 | 101,19 | 7,9500 | 0,770 | 1,52184E-06 | Water | Test |
| Triethylamine | Tertiary amine | 121-44-8 | 101,19 | 57,0700 | 0,726 | 1,43488E-06 | Water | Test |
| Butylamine | Primary amine | 109-73-9 | 73,14 | 92,9000 | 0,740 | 2,02354E-06 | Water | Test |
| Benzaldehyde | Aromatic aldehyde | 100-52-7 | 106,12 | 1,2700 | 1,044 | 1,96751E-06 | Ethanol | Test |
| Beta citronellol | Monoterpene | 106-22-9 | 157,27 | 0,0200 | 0,857 | 1,08985E-06 | Ethanol | Test |
| Formic acid | Carboxylic acid | 64-18-6 | 46,03 | 36,4770 | 1,220 | 5,30147E-06 | Water | Test |
| Acetic acid | Carboxylic acid | 64-19-7 | 60,05 | 15,7000 | 1,049 | 3,49364E-06 | Water | Test IR64a |
| Propionic acid | Carboxylic acid | 79-09-4 | 74,08 | 3,5300 | 0,988 | 2,66734E-06 | Water | Test IR64a |
| Butanoic acid | Carboxylic acid | 107-92-6 | 88,11 | 1,6500 | 1,135 | 2,57644E-06 | Water | Test IR64a |
| Hexanoic acid | Carboxylic acid | 142-62-1 | 116,16 | 0,0435 | 0,929 | 1,59952E-06 | Water | Test |
| Octanoic acid | Carboxylic acid | 124-07-2 | 144,21 | 0,0037 | 0,91 | 1,26201E-06 | Water | Test |
| Water | Oxygen hydride | 7732-18-5 | 18,02 | 24,475 | 0,997 | 1,10686E-05 | Water | IR64a (Solvent) |
| Ethanol | Primary alcohol | 64-17-5 | 40,07 | 59,3000 | 0,790 | 3,9432E-06 | Ethanol | Solvent |

We next compared the odor responses to amines conferred by CpomIR41a1 expression by generating dose-response curves. Mutant ac4 sensilla expressing CpomIR41a1 flies responded to putrescine, spermidine, and hexylamine that we observed as a common activator (Supplementary data file 1) in a dose-dependent manner, and the pharmacological parameters are indicated in Table 2. Our results suggest that CpomIR41a1 is more sensitive to spermidine than putrescine (Figure 2A), as reflected by the EC50 values. Overall, our findings provide a successful example of IR deorphanization using co-expression of a heterologous IR alongside a native IR.

**Figure 2.**
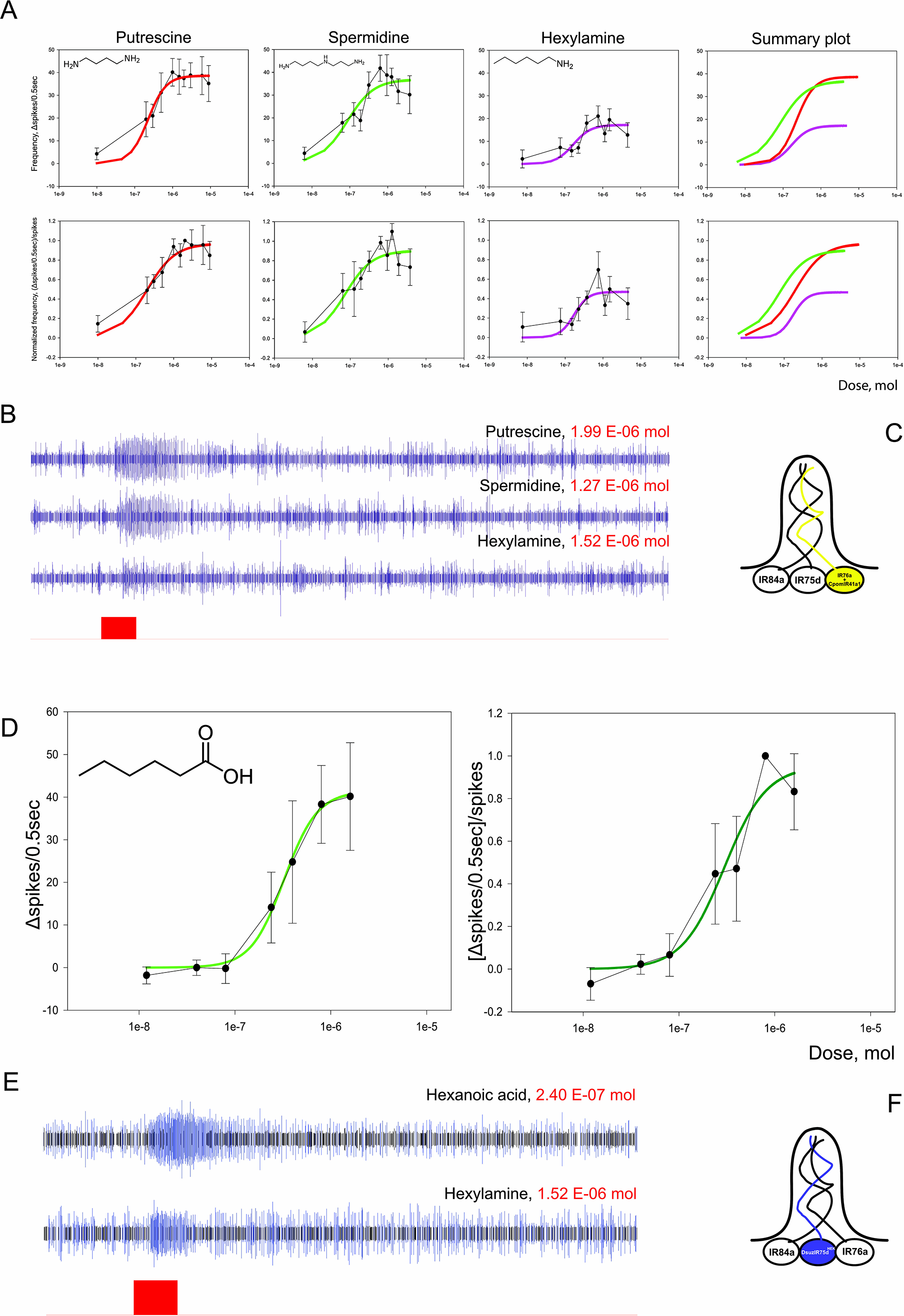
Dose-response characteristics of ac4 sensilla from transgenic lines expressing *CpomIR41a1* and *DsuzIR75d^HEK^*. (**A**) Dose-response characteristics of ac4 sensilla recorded from antennae of *w;pIR76a-Gal4;pUAS-CpomIR41a1* fly lines to putrescine, spermidine and hexylamine. The ac4 sensilla containing IR76a-neurons co-expressing CpomIR41a1 generated spiking in response to these compounds. The effects were concentration dependent and reversible. Left: graphs of concentration dependences of the compounds expressed as a function of spike frequency (Δspikes/0.5sec, above) or normalized frequency [(Δspikes/0.5sec)/spikes, below]. Spike frequency was normalized on the effect to saturating doses of putrescine (DOSE 20: 1.99 E-06 mol; Supplementary data file 3). Error bars represent standard error of means. Data were fit with the Hill equation (solid lines). Right: summary plots; different colors depicts different compounds (putrescine, red; spermidine, green; hexylamine, magenta). (**B**) Spike trains of ac4 generated by DOSE 20 of the specific stimuli (putrescine: 1.99 E-06 mol; spermidine: 1.27 E-06 mol; hexylamine: 1.61 E-06 mol; insect replicate n. 4 from Supplementary data file 1 (see sheet named *w;pIR76a-Gal4;pUASCpomIR41a1*). Red bar: stimulus. (**C**) Schematic representation of ac4-sensilla of transgenic *D. melanogaster* co-expressing CpomIR41a1 in IR76a-neurons. (**D**) Dose-response characteristics of ac4 sensilla recorded from antennae of *w;pIR75d-Gal4/pUAS-DsuzIR75d^HEK^;IR75d^KO^*fly lines to hexanoic acid expressed as a function of spike frequency (Δspikes/0.5sec, left) or normalized frequency [(Δspikes/0.5sec)/spikes, right]. The ac4 sensilla containing IR75d-neurons expressing DsuzIR75d generated spiking in response to application of hexanoic acid. Spike frequency was normalized on the effect to saturating DOSES 10 (8.00 E-07 mol) or 20 (1.6 E-06 mol) depending from the replicate (Supplementary data file 3 – sheet DsuzIR75d). (**E**) Spike trains of ac4 generated by DOSE 3 of hexanoic acid (insect replicate n. 2 from Supplementary data file 3 – sheet *DsuzIR75d*) and DOSE 20 of hexylamine (insect replicate n. 8 from Supplementary data file 1 – sheet *w;UASDsuzIR75d-IR75dGal4;IR75KO*). Red bar: stimulus. (**F**) Schematic representation of ac4-sensilla of transgenic *D. melanogaster* replacing *DsuzIR75d^HEK^* expression to *IR75d* in its specific neurons. Colors have been adopted as shown in Figure 1.

**Table 2.** pharmacological parameters. The table reports potency (EC50), Hill coefficients and maximal effect (Fmax) of ligands from Table 1 found as the main activators for CpomIR41a1 (putrescine, spermidine, hexylamine) and for DsuzIR75d (hexanoic acid) when tested in transgenic *Drosophila*. Values compare raw (from Spikes/0.5 sec) and normalized values ([Spikes/0.5 sec]/spikes). Respective saturating doses are indicated below.

| Normalization | Parameters | CpomIR41a |  |  | DsuzIR75d |
| --- | --- | --- | --- | --- | --- |
|  |  | Putrescine | Spermidine | Hexylamine | Hexanoic acid |
| Not-normalized | EC50 (mol) | (2.23 ± 0.293) E-07 | (8.49 ± 3.27) E-08 | (1.67 ± 0.689) E-07 | (3.26 ± 0.196) E-07 |
|  | Hill coeff. | 1.833 ± 0.6107 | 1.235 ± 0.7201 | 1.864 ± 1.505 | 2.501 ± 0.3859 |
|  | Fmax (spks/0.5s) | 38.62 ± 1.593 | 36.84 ± 4.115 | 17.21 ± 2.849 | 41.38 ± 1.667 |
| Normalized | EC50 (mol) | (1.92 ± 0.457) E-07 | (7.16 ± 2.87) E-08 | (1.73 ± 0.691) E-07 | (2.98 ± 0.892) E-07 |
|  | Hill coeff. | 1.15 ± 0.443 | 1.201 ± 0.683 | 2.311 ± 2.174 | 1.973 ± 1.109 |
|  | Fmax [(spks/0.5s)/spks] | 0.9699 ± 0.06589 | 0.9021 ± 0.09377 | 0.47 ± 0.08032 | 0.9527 ± 0.1688 |
| Saturation (mol) |  | 1.99 E-06 | 6.37 E-07 | 7.57 E-07 | [0.80 ; 1.60] E-06 |

### Functional characterization of D. suzukii IR75d heterologously expressed in IR25a-based “ionotropic receptor decoder” of D. melanogaster

The *D. suzukii* DsuzIR75d was heterologously expressed in the OSN of a transgenic *D. melanogaster* that was lacking the expression of its own DmelIR75d but expressed the co-receptor DmelIR25a. DmelIR75d is known for being activated by pyrrolidine, as well as DmelIR76a (Silbering et al., 2011). Consequently, SSRs from ac4 sensilla always showed a response to this ligand, even in transgenic lines that did not express DmelIR75d (Figure 1). However, when DsuzIR75d was expressed in *D. melanogaster* ac4 sensilla, these neurons became sensitive to stimulation to hexanoic acid (*p* = 0.020) (Figure 1B, Supplementary data file 1), a stimulus that did not activate control ac4 neurons. Further characterization showed that mutant ac4 sensilla expressing DsuzIR75d responded to hexanoic acid in a dose-dependent manner (Figure 2D) pointing that this is an agonist for DsuzIR75d. Additionally, expression of DsuzIR75d reduced the firing frequency of ac4 when exposed to triethylamine (p = 0.011) and abolished butylamine-evoked responses (*p* = 0.436), suggesting that the transgenic expression somewhat interfere with the neuronal response of ac4 sensilla. Such inhibitory effects were also tested in dose response but showed that although constantly inhibitory, there were not exhibiting dose-response characteristics (Supplementary Figure S1).

This result demonstrates that the use of a novel“ionotropic receptor decoder” based on a neuron expressing the co-receptor DmelIR25a but lacking its native receptor is suitable for IR characterization from non-model species.

### Functional characterization of D. suzukii heterologously expressed in IR8a-based “ionotropic receptor decoder” of D. melanogaster

To deorphanize the IR64a tuning subunit from *D. suzukii* (DsuzIR64a), we employed a “ionotropic receptor decoder’’ similar to those used for DsuzIR75d. In this case, we drove the expression of DsuzIR64a in the OSN that lacks the expression of native DmelIR84a but still expresses the co-receptor DmelIR8a (Figure 1C). This strategy has been already proved to be useful in the deorphanization of IRs from *D. sechellia* (Prieto-Godino 2016, 2017, 2021). This choice was based on the fact that in *D. melanogaster*, the IR64a orthologue, DmelIR64a, is dependent on the presence of the co-receptor DmelIR8a (Ai et al., 2010; 2013). Hence, we hypothesized that the conserved orthologous from *D. suzukii* would also require an IR8a co-receptor for proper functioning. The results showed that there were no novel odor-evoked responses in mutant ac4 sensilla expressing DsuzIR64a compared to control ac4. Specifically, we did not observe any response to the ligands known to activate the orthologue DmelIR64a, which senses carboxylic acids (Ai et al., 2010; 2013). This suggests that under our experimental conditions, the expressed IR tuning subunit was either not functional or had a shift in its binding affinity towards ligands not present in the tested panel. However, expression of DsuzIR64a interfered with the normal firing activity of ac4, reducing the spike frequencies of mutant ac4 sensilla when exposed to pyrrolidine (p = 0.174), dimethylamine (p = 0.462), ammonium hydroxide (p = 0.175) compared to control ac4 (Figure 1C; Supplementary data file 1). Interestingly, although at the limit of significance, reduction in firing activity of ac4 was observed also for butylamine (p = 0.055).

In a parallel set of experiments conducted by expressing a truncated isoform of the *C. pomonella* orthologue CpomIR64a lacking the transmembrane region M3 (UniProt acc. num. A0A0V0J232), we observed the same reduction in firing activity to the above mentioned stimuli (pyrrolidine, p = 0.278; dimethylamine, p = 0.816; ammonium hydroxide, p = 0.058), indicating that heterologous expression of novel IR subunits in the OSN lacking IR84a may interfere with the native neuronal activity.

### Fluorescent in situ hybridization and structural analysis of DsuzIR64a

Since DsuzIR64a was not activated by the carboxylic acids known to activate the orthologous DmelIR64a, we further explored the possible molecular basis of this evolutionary switching. First of all, we checked if DsuzIR64a was expressed at the same location of DmelIR64a, testing a parallel set of antennae for positive (Orco) and negative (IR62a) controls, as done in previous projects (Cattaneo et al., 2022). DmelIR64a is one of the *D. melanogaster* antennal IRs expressed in the OSNs surrounding the third saccular chamber (Benton et al., 2009; Ai et al., 2010). Since the antennal organization of *D. suzukii* resembles that of *D. melanogaster*, we asked if this IR was expressed in the same antennal region also in this species. Results from fluorescent *in situ* hybridization (FISH) indicated that ∼26 OSNs in proximity to the sacculus express DsuzIR64a (Figure 3 A). There were no differences between males and females (Figure 3B) (Mann–Whitney-U-test: U = 38; p = 0.322, Supplementary data file 2).

**Figure 3.**
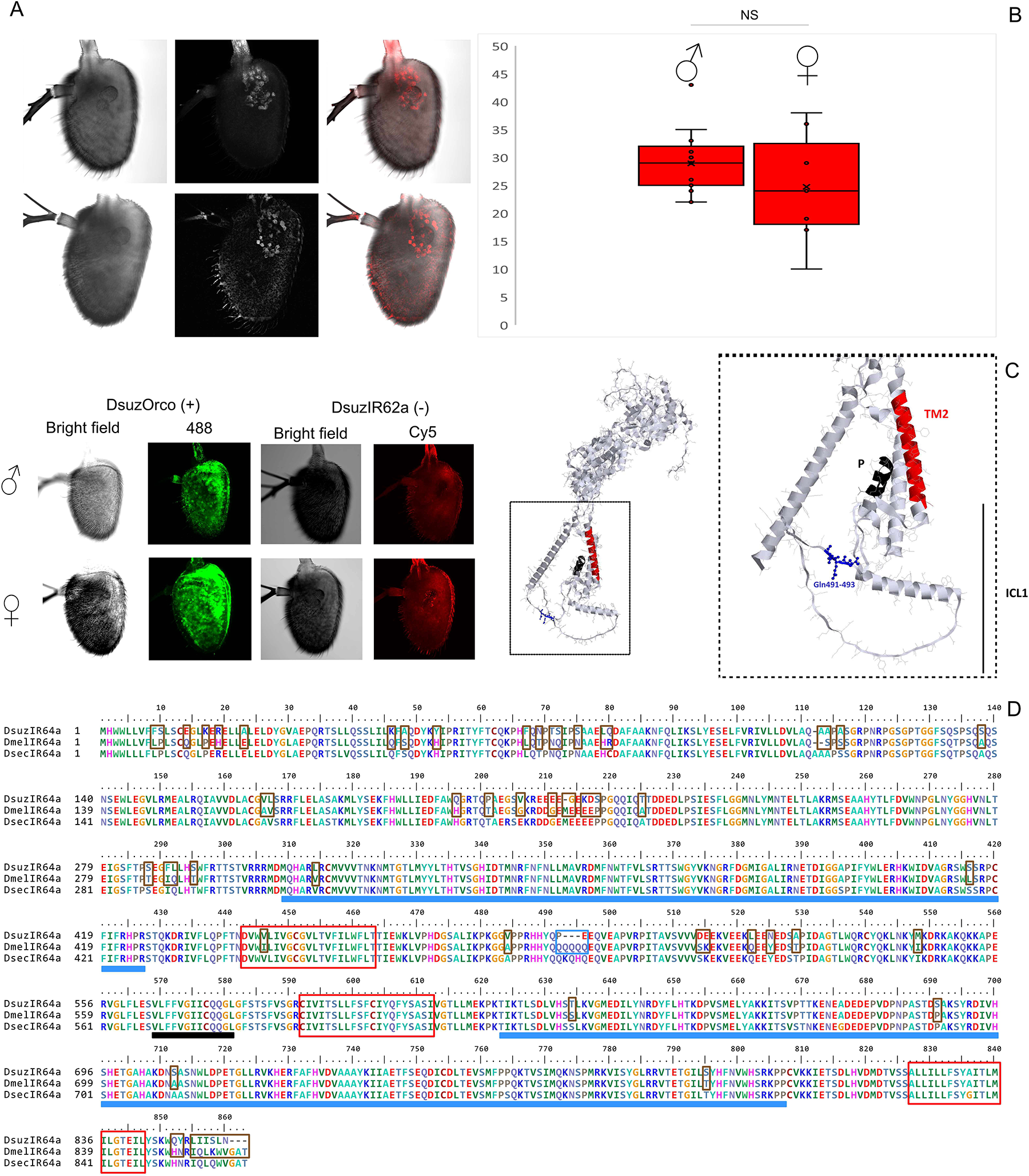
Fluorescent *in situ* hybridization of DsuzIR64a and structural analysis. (**A**) Fluorescent *in situ* hybridization of the DsuzIR64a subunit comparing male (♂) and female (♀) antennae from bright field (left), Cy5 and merged channels (N_male_ = 15, N_female_ = 7). Specifically, the male antenna shown in this figure represents replicate 3, while the female antenna represents replicate 6 (Supplementary data file 2). Control experiments have been conducted in parallel with FISH for DsuzIR64a testing expression of Orco (positive control) DsuzIR62a (negative control). (**B**) Boxplots generated upon analysis of neuronal counting (Supplementary data file 2). NS: no significant differences in terms of the number of neurons were validated for the staining of IR64a transcripts (MWU-test: *p* = 0.32218; ::J = 0.05; U = 38). (**C**) Left: protein model of DmelIR64a generated by RasTop. Right: magnified view of the ICL-1 of DmelIR64a. Blue: glutamine residues absent in the subunit of *D. suzukii*. Black: pore loop; Red: transmembrane domain M1, as in Benton et al., (2009). (**D**) Polypeptide sequence alignment of IR64a between *D. suzukii*, *D. melanogaster* and *D. sechellia*. Amino acid substitutions, insertions and deletion between *D. suzukii* and *D. melanogaster* are represented as brown opened squares, while the deletion of the glutamine residues indicated in C is represented as a blue opened square. Structural domains are represented as filled blue squares (LBD, S1 and S2), a filled black square (pore loop) and opened red squares (transmembrane domains).

Secondly, we carried out a sequence alignment analysis of *D. melanogaster*, *D. sechellia* and *D. suzukii* IR64a (Figure 3, Supplementary Figure S2) to identify the likely molecular basis of subunit specificity switch. Examination of the identity of residues aligned with DsuzIR64a revealed a high conservation across the three drosophilid species although several substitutions, including one insertion and three deletions, were exclusively present in the *D. suzukii* IR64a orthologue copy. In particular, the most evident deletion is located at the Q490 position of the DmelIR64a orthologue, in which three consecutive glutamines are missing in the DsuzIR64a subunit, which folding unveiled their positioning within the half of the intracellular loop 1 (Figure 3C). However, none of these insertions and deletions are located in the ligand binding domain (LBD) of DsuzIR64a, which presents only six amino acid substitutions (Figure 3D, Supplementary Figure S2), distributed as two in the S1 subunit and four in the S2 subunit.

### Structural analysis of acid sensing IRs

Structural analysis was performed in search of specific amino acid residues from the LBD to explain our evidence of hexanoic acid binding for DsuzIR75d and absence of acid binding for DsuzIR64a (Figure 1). To identify these residues, we first conducted a polypeptide sequence alignment (Supplementary Figure S2): since there are no studies investigating the structures of IR75d and IR64a subunits, we advance hypotheses based on the most recent studies on the IR75b and IR75a acid sensors (Prieto-Godino et al., 2017, 2021). Analysis was performed on predicted IR-structures from *D. melanogaster*, *D. sechellia* and *C. pomonella* because of the given absence of deposited structures from *D. suzukii* (Figure 4).

**Figure 4.**
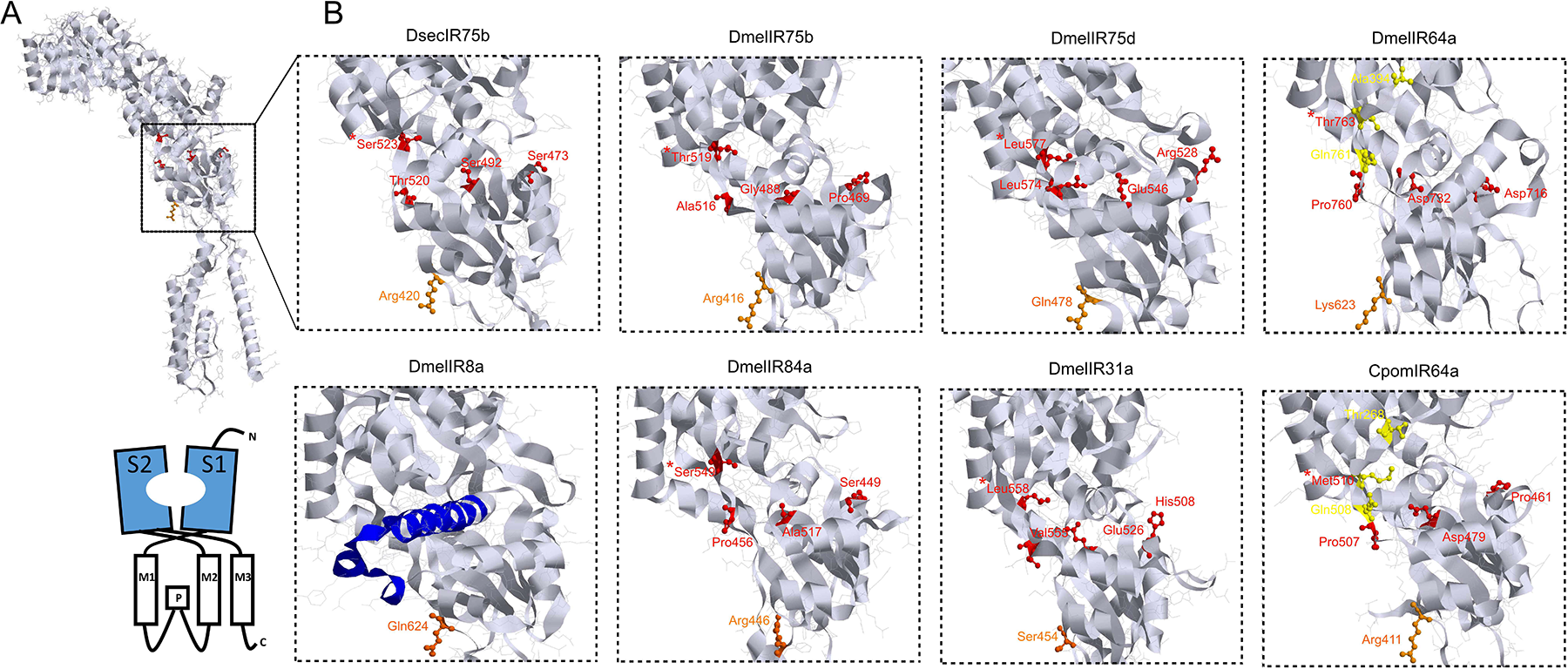
Structural analysis. (**A**) Protein model of DsecIR75b generated by RasTop. Below: cartoon of the domain organization of IR75b (**B**): magnified view of the ligand-binding domain (LBD) for DsecIR75b, DmelIR75b, DmelIR75d, DmelIR64a, DmelIR8a, DmelIR84a, DmelIR31a and CpomIR64a. Red residues: amino acid residues of the LBD involved in hexanoic acid binding of IR75b, based on evidence from Prieto-Godino et al., (2017). Yellow residues: amino acid residues of the LBD involved in C1-C6 acid binding of IR75a, based on evidence from Prieto-Godino et al., (2021). Red/yellow asterisked residue: residue corresponding to the DsecIR75b Ser523 (Prieto-Godino et al., 2017) and the Dmel/DsecIR75a Phe/Leu538 (Prieto-Godino et al., 2021). Orange residue: conserved positively charged polar residue among the acidic sensing IRs. Note: this residue is substituted to serine in DmelIR31a (Ser454) and to glutamine in DmelIR75d (Gln478), DsecIR75d (Gln484) and DmelIR8a (Gln624) (Supplementary Figure S2). Blue: IR8a CREL (Abuin et al., 2019).

By comparing the IR75b sequence of *D. melanogaster* and *D. sechellia*, Prieto-Godino et al., (2017) unveiled eleven key substitutions within the ligand binding domain (LBD), among which, four (Ser473, Ser492, Thr520 and Ser523) lie within the ligand-binding pocket, where the presence of Ser523 is sufficient for binding hexanoic acid. Observing that for DmelIR75d (Figure 4) and DsuzIR75d (Supplementary Figure S2) the key LBD-residues reported in DsecIR75d by Prieto-Godino et al., (2017), which are responsible to hexanoic acid binding, are not conserved, we may exclude their possible involvement in binding to this ligand for the IR75d orthologues. However, although theoretically, we may rather assume that for *D. suzukii* other residues of the IR75d LBD may interact with this ligand. Interestingly, all acid sensing IRs except for DmelIR31a share a positively charged amino acid at the beginning of the S2 (Figure 4, Supplementary Figure S2), which corresponds to Arg416 in DsuzIR75d. In support to a possible involvement of this residue in hexanoic acid binding capacities for the *D. suzukii* orthologue (Figure 1B), IR75d from the *D. melanogaster* group, which do not bind acids (Silbering et al., 2011), presents a polar uncharged amino acid (Gln478), which is the sole non-conserved substitution when compared with the LBD of the orthologue of *D. suzukii*.

Analyzing the structure of DmelIR64a, we noticed that despite the substitution Thr763 is structurally analogous to Ser523 from DsecIR75b (Prieto-Godino et al., 2017), its identity with the residue from the *D. melanogaster* IR75b orthologue (Thr519) that does not sense hexanoic acid, excludes the possible involvement of this residues in hexanoic acid binding for DmelIR64a (Ai et al., 2010). Similar evidence may be expected for DsuzIR64a, which conserves the same residues of DmelIR64a (Supplementary Figure S1). Out of the three key residues commuting different C1-to-C6 acid sensing between the IR75a orthologues of *D. melanogaster* and *D. sechellia* (T289S, Q536K and F538L; Prieto-Godino et al., 2021) two of these residues in DmelIR64a are not conserved (A394 rather than T289S and T763 rather than F538L, Figure 4). For this reason, the documented binding of C1-C6 acid ligands for DmelIR64a (Ai et al., 2010, 2013) may involve other residues within the LBD. Similar evidence is expected for DsuzIR64a, which conserves the same residues of DmelIR64a (Supplementary Figure S1). While further studies may address the identification of possible residues influencing acid binding within the IR64a sequences, we did not observe outstanding differences between the LBDs of *D. melanogaster* and *D. suzukii* IR64as (Figure 3D) to advance explanation of lack of acid binding from the latter (Figure 1C). However, polypeptide sequence alignment unveiled several amino acid substitutions, insertion and deletions between these two subunits (Supplementary Figure S2, Figure 3D). Although hypothetically, such differences may hide the incapacity of binding acid ligands from the orthologue of *D. suzukii*.

### Heterologous expression of insect IRs in HEK293T cells

We next explored the utility of an *ex vivo* system, specifically the HEK293T system, for heterologous expression and functional characterization of insect IRs. Initially, we optimized the expression protocol using *D. suzukii* DsuzIR8a. To confirm correct expression of this transgene, we stained co-transfected HEK293T cells with an anti *D. melanogaster* IR8a conjugated with Alexa488 dye. This antibody is capable of recognizing DsuzIR8a due to the high sequence conservation. As shown in Figure 5A, the majority of co-transfected HEK293T cells carrying the pcDNA-3.1-DsuzIR8a showed the Alexa488 signal. Furthermore, the signal was localized around the cytoplasm expressing the EBFP2 blue fluorescent protein (EBFP), which served as transfection control. This indicates proper expression of DsuzIR8a in the cell membrane, leading us to conclude that HEK293T cells are a suitable system for accurate IR expression.

**Figure 5.**
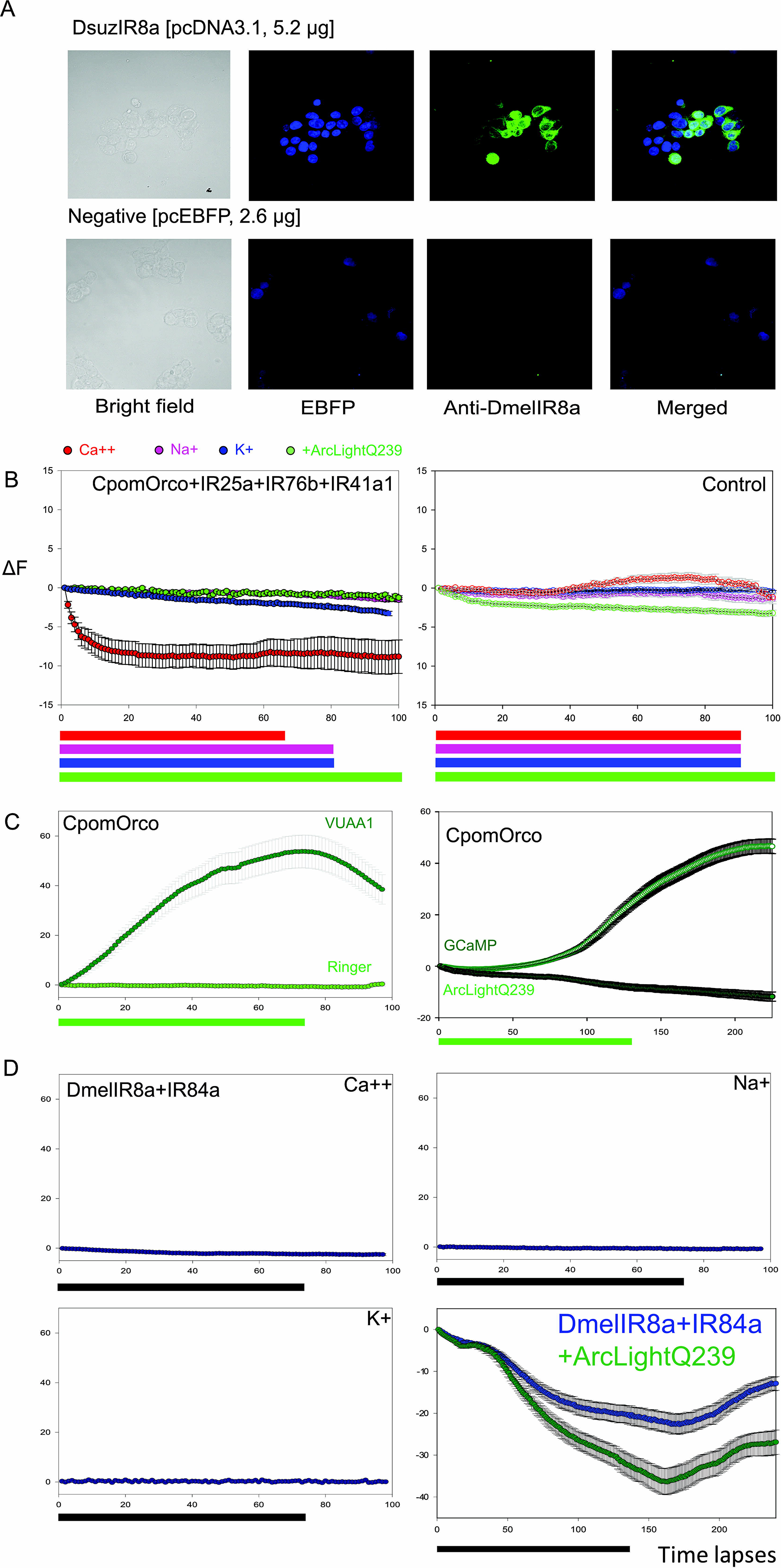
Heterologous expression of insect IRs in HEK293T cells. (**A**) Optimized transfection of HEK293T cells expressing *D. suzukii* IR8a. Immunohistochemistry was performed as in our previous studies (Cattaneo et al., 2017) but combining a Guinea pig polyclonal Anti-DmelIR8a primary antibody [antigen: GLYARRNLGASDSHSGYME (C-terminus)] with an Alexa488 anti-Guinea pig goat-polyclonal. (**B**) Left: fluorescent effect of HEK293T cells stimulated pyridine 1.0-2.0 mM, when co-transfected with CpomOrco+IR25a+IR76b+IR41a1 (N = [37-77]). Right: effects from the respective controls (N = [65-72]), measuring selectivity to ions [Ca^++^ (GCaMP, red); Na^+^ (NaTRIUM Green-2 AM, magenta), K^+^ (ION Potassium Green-4 AM, blue)] and voltage (+ArclightQ239, green). Stimulus duration (at least 65% of the experiment) varies for different lengths of the experiments (colored bars). (**C**) Left: positive control testing the Orco-ligand VUAA1 (green bar) at 500 μM on HEK293T cells expressing CpomOrco, comparing stimulation with ringer buffer+DMSO (N = 53). Right: optimization of an ArcLight-based fluorimetric method: CpomOrco+GCaMP (N = 129) versus CpomOrco+ArcLightQ239 (N = 74); green bar: VUAA1 500 μM. Note: decrement in fluorescence for ArcLightQ239 indicates access of cations into the plasma membrane. (**D**) Functional characterization attempts testing 1.0 mM phenylacetic acid (black bar) on HEK293T cells expressing IR8a of *D. melanogaster* with IR84a (DmelIR8a+IR84a). Response of DmelIR8a+IR84a to the IR84a-ligand phenylacetic acid (black bar) was compared for selectivity to Ca^++^ (N = 115), Na^+^ (N = 89) and K^+^ (N = 54) using the same indicators from (B). Bottom right: comparison of the fluorescent effect of HEK293T cells co-transfected with DmelIR8a+IR84a+ArcLightQ239 (blue, N = 67) with HEK293T cells transfected with the sole ArcLightQ239 (green, N = 59) when stimulated with 2.0 mM phenylacetic acid (black bar). Vertical bars in plots: standard errors.

Next, we co-transfected HEK293T cells with an expression plasmid carrying *CpomIR41a1*, alongside the broadly-expressed co-receptors *C. pomonella* IR25a (CpomIR25a) and IR76b (CpomIR76b), which are required for its function. Additionally, a plasmid carrying the odorant receptor co-receptor Orco (CpomOrco) was also co-transfected as a positive control. We perfused the co-transfected cells with pyridine, a ligand we had previously demonstrated to activate CpomIR41a1 in *in vivo* expression experiments (Figure 1A). We observed an overall decrease of fluorescence when testing cation permeability and voltage. Control experiments also revealed an apparent increment in fluorescence associated with calcium and a decrement in voltage-associated fluorescence, suggesting that the expression of the combination IR41a-IR25a-IR76b in HEK293T cells did not produce a specific response (Figure 5B). However, when we perfused VUAA1, an activator of CpomOrco (Cattaneo et al., 2017), which was co-transfected with the plasmid carrying the three IRs, a clear response was observed when monitoring both calcium and voltage (Figure 5C). This indicates that the system was functional and the lack of response from the IR combination was likely due to other effects.

Lastly, we attempted to use HEK293T cells to deorphanize a tuning IR which requires the IR8a co-receptor instead of IR25a. We specifically selected the *D. melanogaster* DmelIR84a since its main agonist phenylacetic acid is already known (Grosjean et al., 2011). When we perfused phenylacetic acid we did not observe any discernible effects (Figure 1 D). However, the voltage indicator ArcLightQ239 unveiled a reduction in the amplitude of fluorescent variation, possibly indicating the DmelIR8a+DmelIR84a activation by phenylacetic acid. Nevertheless, we observed the same effect when ArcLightQ293 was expressed alone, suggesting that this decrease in fluorescence was likely an artifact. In conclusion, our results indicate that, under our experimental conditions, HEK293T cells are not suitable for the functional characterization of IRs.

## Discussion

In-depth studies have been conducted on IRs in *Drosophila* and a few mosquito species. However, there is still a significant lack of information regarding the role of these receptors in odor perception in other insect orders. This knowledge gap primarily stems from the limited availability of genetic tools in non-model species, which hampers the functional characterization of tuning subunits outside of *Drosophila*. In this study, we expanded the deorphanization strategy of the “ionotropic receptor decoder” from using an OSN that expresses the co-receptor DmelIR8a (Prieto Godino et al., 2016; 2017; 2021) to other OSNs expressing the co-receptor DmelIR25a or the combination of DmelIR25a and DmelIR76a. Furthermore, we demonstrated for the first time the effectiveness of this strategy in characterizing IR tuning subunits from fruit flies outside the *D. melanogaster* species subgroup, as well as from distantly related orders such as Lepidoptera.

Evolutionary studies have revealed that IR subunits diverge in terms of gene numbers and coding sequences within and across species, while the co-receptors remain highly conserved (Croset et al., 2010, Yin et al., 2021). This suggests that the assembly and function of heteromeric complexes are likely to be similar across different lineages. Indeed, transgenic expression of DsuzIR75d in *D. melanogaster* DmelIR25a-expressing OSNs led to the formation of a functional heteromeric complex, as did the transgenic expression of CpomIR41a1 in another OSN expressing both DmelIR25a and DmelIR76b. This demonstrates the broad conservation of the IR-based olfactory system across insects, and opens up avenues for the widespread use of this technique in IR deorphanization. This approach may parallel the extensive use of the *Drosophila* “empty neuron system” for deorphanizing ORs from non-model species. The *Drosophila* “empty neuron system” has been employed not only for functionally characterizing dipteran ORs outside the Drosophilidae lineage (Chahda et al., 2019; Carey et al., 2010) but also for species from Lepidoptera (Kurtovic et al., 2007; Llopiz-Gimenez et al., 2021; de Fouchier et al., 2017; Cattaneo 2018; Syed et al., 2010; Syed et al., 2006) and Orthoptera (You et al., 2016; Chang et al., 2022). This growing body of data has enabled comparative studies across families and orders, which has enhanced our comprehension of the species adaptations driving ecological interactions in the real world (Guo et al., 2021).

The “ionotropic receptor decoder” system offers several advantages over the traditional *ex vivo* approach, such as heterologous expression in *Xenopus* oocytes, which has been used in a limited number of non-model species for IR deorphanization (Xu et al., 2014; Ray et al., 2023; Pitts et al., 2017; Shan et al., 2019; Hou et al., 2022). One of the main advantages of using *an in vivo* system like the “ionotropic receptor decoder” is that it closely mimics the native olfactory sensillum environment, ensuring a high-fidelity receptive field for the expressed receptor. Additionally, the vapor-phase odor delivery in the empty-neuron technique provides a more realistic physicochemical environment compared to water-phase odor delivery required by *in vitro* systems. Altogether, these differences contribute to the increased sensitivity of *in vivo* deorphanization compared to the oocyte expression system (Gupta et al., 2022; Wang et al., 2016). However, there are also some disadvantages to using an *in vivo* system. The generation of transgenic fly lines and the subsequent electrophysiological recording experiments can be time-consuming and may not be suitable for high-throughput screenings. In our experiments, we demonstrated that the deorphanization strategy based on transgenic expression of IRs in *D. melanogaster* OSNs can be achieved using both empty OSNs and OSNs that already express native IR subunits (Figure 1, 2). This flexibility in expression methods facilitates the use of this technique, as it does not necessarily require the creation of transgenic *D. melanogaster* IR knockout lines. However, it is important to exercise caution and implement appropriate controls, as the firing profile of the native IR subunits may potentially mask the response of the transgenic IRs (Figure 1).

Insect IR repertoires consist of two subfamilies: conserved antennal IRs (A-IRs) and the species-specific divergent IRs (D-IRs) (Croset et al., 2010). Antennal IRs are conserved in sequence and expression patterns across different insect orders. For example, certain A-IRs found in Drosophilidae genomes are also present in Lepidoptera, Coleoptera, and Phthiraptera species (Yin et al., 2021; Croset et al., 2010). IR41a is one such example, since it exhibits a one-to-one copy in all analyzed lepidopteran and dipteran genomes, with additional copies found in specific mosquito and lepidopteran species (Pitts et al., 2017; Yin et al., 2021; Croset et al., 2010), including *C. pomonella*, where it is present in two putative paralogues (Walker et al., 2016). In *D. melanogaster* DmelIR41a forms a heteromeric complex with DmelIR25a and DmelIR76b, responding to polyamine ligands like pyridine, pyrrolidine, putrescine, cadaverine and spermidine (Silbering et al., 2011; Hussain et al., 2016). A similar activation pattern is observed in the mosquito *A. gambiae* (Pitts et al., 2017). Our study extends this functional conservation beyond the Diptera order, as the mutant ac4 expressing CpomIR41a1, DmelIR25a, and DmelIR76b exhibited firing activity when stimulated with pyridine, putrescine, cadaverine, and spermidine. However, stimulation by pyrrolidine could not be confirmed due to spontaneous firing caused by native DmelIR76a and DmelIR75d expressions (Silbering et al., 2011).

Another broadly conserved A-IR is IR75d, which belongs to the IR75 clade (Croset et al., 2010; Silbering et al., 2011, Yin et al, 2021; Huo et al., 2022). In *D. melanogaster*, this clade has four paralogs. Three of them, DmelIR75a, DmelIR75b and DmelIR75c are activated by C2-C6 carboxylic acids (Abuin et al., 2011; Silbering et al., 2011; Gorter et al., 2016; Prieto-Godino et al., 2016, 2017, 2021) whereas DmelIR75d agonist is activated by the polyamine pyrrolidine but not by carboxylic acids (Silbering et al., 2011; Prieto Godino et al., 2017). Of these four IR75 subunits, only IR75d maintains a clear one-to-one orthologous relationship across dipteran and lepidopteran lineages, and these latter have additional copies of IR75 isoforms that are not orthologous to DmelIR75a, DmelIR75b, and DmelIR75c (Croset et al., 2010; Yin et al., 2021; Hou et al., 2022). Several of these IR75 subunits have been functionally characterized, primarily showing tuning to carboxylic acids. For instance, AaegIR75k1 and AaegIR75k3 from *Aedes aegypti*, as well as AalbIR75e from *Aedes albopictus*, respond to C7-C9 carboxylic acids (Ray et al, 2023), and the agonist of *A. gambiae* AgIR75k are C6-C10 carboxylic acids (Pitts et al., 2017). Lepidopteran IRs AsegIR75p.1 and AsegIR75q.1 from *Agrotis segetum* also respond to medium-chain fatty acids, with hexanoic acid being the most potent agonist for AsegIR75p.1 (Huo et al., 2022). In our screening, we were unable to evaluate the conservation of pyrrolidine-evoked responses in ac4 sensilla, as the firing activity by DmelIR76a-expressing OSNs masked the response of DsuzIR75d (Abuin et al., 2011; Silbering et al., 2011, Vulpe and Menuz 2021). However, further screening identified hexanoic acid as an agonist for DsuzIR75d, suggesting a conserved functional specialization for carboxylic acids that originated in the ancestral IR75 receptor and that has been lost in DmelIR75d. In support to this hypothesis, a conserved arginine proximal to the end of the LBD-S2 domain, renowned as one of the main agonist-binding residues characteristic of iGluRs that is globally conserved in acid-sensing IRs, but not in non-acid-sensing IRs (Armstrong and Gouaux 2000), is also present in the orthologue of *D. melanogaster* (Supplementary Figure S2).

Finally, our screening identified a novel agonist for *D. melanogaster* ac4 sensilla: hexylamine. This compound was not included in the stimuli screened thus far against this sensillum (Benton et al., 2009; Silbering et al, 2011; Grosjean et al., 2011; Abuin et al., 2011; Hueston et al., 2016; Abuin et al., 2019; Prieto-Godino et al., 2016; 2017; 2021). The discovery of hexylamine as an agonist aligns with previous suggestions that sensing of hexylamine, which repels *D. melanogaster*, is mediated by OSNs other than those expressing DmelIR92a and housed in ac1 sensilla. This is supported by the negligible effects on hexylamine-driven behavioral response when DmelIR92a-expressing neurons are inactivated (Min et al., 2013). It would be intriguing to understand which IR tuning subunit expressed in ac4 sensilla is responsible for hexylamine sensing. Our results showed that ac4 sensilla from both knock-out lines, lacking the expression of DmelIR84a and DmelIR75d, exhibited a firing response to hexylamine stimulus similar to ac4 sensilla from the control line. This observation rules out the possibility that either of these two IR subunits is necessary for sensing hexylamine. However, it leaves open the possibility that both subunits contribute redundantly to the response to this volatile compound. Further studies are needed to understand the contribution of these subunits, as well as DmelIR76a, in sensing hexylamine.

When we expressed DsuzIR64a in the “ionotropic receptor decoder” system previously used for deorphanization of IR subunits from *D. melanogaster* and *D. sechellia* (Prieto-Godino et al., 2016; 2017; 2021, Abuin et al., 2011, Grosjean et al., 2011) there was no new firing activity compared to the control knock-out line. This suggests that either the IR64a-transgene expression was not functional or the specific activating ligands were not present in the odor panel used for screening. In support to this second hypothesis we showed that, despite the ligand binding domain (LBD) of DmelIR64a and DsuzIR64a subunits are mostly identical, several substitutions, insertions and deletions were identified for the rest of their sequences (Supplementary Figure S2, Figure 3). Among them, we identified a deletion of three consecutive glutamines within the intracellular loop 1 of the DsuzIR64a subunit (Figure 3C). It is known that in other olfactory cation channels in insects, small changes in the amino acid sequence of intracellular loops may influence the ligand binding ability and its pharmacology (Bobkov et al., 2021; Corcoran et al., 2018; Hopf et al., 2015; Turner et al., 2014). Future comparative studies may address investigation from the various differences that we have observed in the sequence of DsuzIR64a, with the aim to understand their possible influences in ligand binding.

Interestingly, in ac4 sensilla expressing both DsuzIR64a and a truncated CpomIR64a isoform, the firing frequency decreased when stimulated with pyrrolidine, dimethylamine, and ammonium hydroxide compared to the control knockout line, and it decreased also to butylamine when DsuzIR64a was expressed (Figure 1C). This reduction in stimulation may be associated with the expression of heterologous subunits, either functional or not. We are aware that this effect was not observed during previous studies that used the “ionotropic receptor decoder” neuron (Benton et al., 2009, Prieto-Godino et al., 2017, 2021). However, these studies did not test a wide panel of ligands such as in our study but rather focused on effects of specific compounds. A similar phenomenon was also observed in the transgenic line expressing CpomIR41a1, which inhibited the responses to phenylacetaldehyde, dimetylamine and phenylacetic acid. These ligands activate DmelIR84-expressing OSNs (Grosjean et al., 2011; Silbering et al; 2011), which were not the target of CpomIR41a1 transgene expression. Similarly, the expression of DsuzIR75d inhibited ac4 response to butylamine, which was detectable in the corresponding knock-out control line. These observations are in discordance with previous findings that agonists for one OSN can antagonize the activity in another (Silbering et al., 2011), since we recovered the same effect also when we expressed the truncated CpomIR64a. Future studies are definitely needed to understand why and how some ectopically or heterologously expressed IR tuning subunits alter the firing responses of other OSNs housed in the same sensillum.

Our proposed comparison of IR75d with IR75b was the most logic from the reported evidence of hexanoic acid binding of the *D. sechellia* orthologue of the latter, in which specific LBD-residues confer response to this ligand (Prieto-Godino et al., 2017). Contrary to our expectations, analysis of the polypeptide sequence and structure of DsuzIR75d (Supplementary Figure S2, Figure 4) unveiled absence of the main S2-residues, which were demonstrated to be essential for hexanoic acid binding in the IR75b orthologue of *D. sechellia* (Prieto-Godino et al., 2017). Interestingly, in comparison to the *D. melanogaster* subgroup, among the few substitutions presented by the IR75d of *D. suzukii* there is a positively charged residue at the beginning of the S2 (Arg411), which is conserved in all the other acid sensing IRs except for DmelIR31a (Supplementary Figure S2). Despite being part of the S2, structural analysis unveiled this residue position external from the LBD (Figure 4), which suggested that any possible involvement of this residue in ligand binding might take place within additional pockets formed by the IR-cation channel. Future studies may focus on this residue to validate the hypothesis of its eventual role in carboxylic acid binding, which raises interesting questions about the functional specialization and evolution of IR75d in switching their sensitivity from acids to polyamines. To investigate further, testing the sensitivity of IR75d to pyrrolidine and hexanoic acid across different Dipterans would shed light on whether the switch to polyamine is specific to *D. melanogaster* or if both responses can coexist for the same IR subunits from other insects.

In this work, we also explored the heterologous expression in HEK cells, which is a single-cell *ex vivo* system such as *Xenopus* oocytes. HEK cells have been largely used in the deorphanization of ORs (Cattaneo et al., 2017; Bobkov et al., 2021, Crava et al., 2022, Miazzi et al., 2019, Grosse-Wilde et al., 2007; 2006) whereas, to our knowledge, this method has not been previously utilized for IR subunit deorphanization. We conducted several experiments using this approach but did not achieve successful deorphanization of IRs, despite observing correct expression of IRs in transfected cells through immunohistochemistry experiments (Figure 5). This may be attributed to several factors. Firstly, the distinct cellular environment of HEK cells compared to OSNs could contribute to the differences in receptor function and signaling. Additionally, the poor surface localization of IR proteins in HEK cells could be a limiting factor, as it has been observed in HEK cells expressing ORs where proteins were retained in intracellular membranes hindering their proper functioning and interaction with ligands (German et al., 2013; Miazzi et al., 2019). These results highlight the challenges associated with the heterologous expression approach in HEK cells for IR deorphanization. Further studies are needed to optimize this method and address the limitations encountered.

### Conclusions

Adding to previous efforts conducted to functionally characterize insect IRs by heterologous expression, our study illustrates that the replacement of tuning IRs in OSNs of *D. melanogaster* with foreign IRs from different insects, as well as the expression of these alongside the native *Drosophila* IRs, leads to the formation of functional heteromeric complexes. For the first time, we have used *D. melanogaster* to deorphanize IR subunits of insects not belonging to its close phylogeny. This achievement not only contributes to the ongoing deorphanization endeavors concerning chemosensors in pests like *C. pomonella* and *D. suzukii* but also holds the potential to guide future investigations into IRs from various other species. Future projects may attempt a transition *in vitro* of this method starting from the pan-neuronal expression of heterologous IRs and the isolation of neurons from transgenic *Drosophila* (Egger et al., 2013, Ui et al., 1994), aimed to direct cell-based approaches to a deeper pharmacological study of these receptors.

Once more, the successful use of *D. melanogaster* proved instrumental in advancing methodologies aimed to study insect chemoreceptors. Our results incorporate the use of transgenic *D. melanogaster* into the toolkit of heterologous methods for *in vivo* functional analysis of IR subunits, spanning from ligand binding to pharmacology.

## Material and methods

### Insects

Transgenic *D. melanogaster* were maintained on a sugar-yeast-cornmeal diet (https://bdsc.indiana.edu/information/recipes/bloomfood.html) at room temperature (25 ± 2°C) and a relative humidity of 50 ± 5 % under 12:12 light: dark photoperiod.

### Amplification and cloning of C. pomonella and D. suzukii IRs

The coding sequences of *C. pomonella* and *D. suzukii* IRs were identified from antennal transcriptome in the frame of three of previous studies (Ramasamy et al., 2016; Walker et al., 2016; Walker et al., 2023). Briefly, part of the total RNA samples used for these studies extracted and purified with a combined approach of TRIzol-based extraction followed by RNeasy® Mini spin column purification (Qiagen, Venlo, Netherlands), was retro-transcribed to cDNA using RT-for-PCR kit (Invitrogen, Life technologies, Grand Island, NY, USA). The complete ORFs encoding CpomIR41a1 was amplified combining forward (5’-CACCATGATAATGCCAAGTAAAC-3’) and reverse (5’-TTATAACACTATTGAAGGTTTGC-3’) primers with upstream *attB*-regions suitable for BP-clonase-recombination (*attB1*: 5′-GGGGACAAGTTTGTACAAAAAAGCAGGCTTAACA-3′; *attB2*: 5′-GGGGACCACTTTGTACAAGAAAGCTGGGT-3′, Gateway Technology, Invitrogen). By same the method, *attB*-regions were integrated also to primers to amplify both the truncated CpomIR64a isoform, UniProt A0A0V0J232 (forward: 5’-CACCATGAACTTAACAACATACAGTG-3’; reverse 5’-TCAAACCTGTAGCTTCTTTAACC-3’), and DsuzIR62a, that we have used to generate synthetic probes for *in situ* negative control experiments, (forward: 5’-CACCATGTTCCTCCAGTTGCTG-3’; reverse 5’-TTAATCCCCGGGGTTGCGGTTAG-3’). The coding sequence of DsuzIR64a, cloned into pDONR221, was obtained upon synthesis (GeneArt Thermo Fischer). Because of the impossibility in amplifying IR75d from *D. suzukii* cDNA and in obtaining a synthetic sequence from GeneArt, we decided to *attB*-combine forward (5’-CACCATGAAGGTCCAGGTGTTCAG-3’) and reverse (5’-TCAGCGCTTCATCTCTTGGTAGC-3’) primers to amplify and clone for *Drosophila* transgenesis the same human HEK-codon-optimized sequence of DsuzIR75d, cloned into pcDNA3.1, that we have used for the expression into HEK293T (from now on, DsuzIR75d^HEK^). For amplification, a temperature program of 98 °C for 5 min followed by 45 cycles of 98 °C for 1 min, Tm of the primer for 1 min and 72°C for 70-90 sec, depending on experiment, with a final elongation step of 68 °C for 7 min. Purified PCR products were then cloned into the pDONR221 plasmid (Invitrogen). For CpomIR41a1, CpomIR64a, DsuzIR75d^HEK^ and DsuzIR64a, cassettes with inserts were then transferred from their cloning plasmids to the destination vector (pUASg-HA.attB, constructed by E. Furger and J. Bischof, kindly provided by the Basler group, Zürich), using the Gateway LR Clonase II kit (Invitrogen). Integrity and orientation of inserts was checked further by Sanger sequencing before using plasmids to generate transgenic *Drosophila*.

### Heterologous expression in D. melanogaster

Transformed lines for pUAS-CpomIR41a1, pUAS-DsuzIR75d^HEK^, pUAS-DsuzIR64a and pUAS-CpomIR64a were generated by Best Gene (Chino Hills, CA, USA) through PhiC31 standard integration either on the 3rd chromosome, injecting BDSC#8622 strains for CpomIR41a1, or on the 2nd chromosome, injecting attp40 strains for DsuzIR75d^HEK^, DsuzIR64a and CpomIR64a. Crossings were performed as shown in Supplementary Figure S3. In brief, we used balancer lines in accordance with procedures already published from our labs (Gonzalez et al., 2016). To generate CpomIR41a1 lines, we used pIR76a-Gal4 parental lines upon balancing (*w;pIR76a-Gal4;TM2/TM6B* – Silbering et al., 2011). To generate DsuzIR75dHEK lines, we combined crossings using a pIR75d-Gal4 line (*w; pIR75d-Gal4;TM2/TM6B* – Silbering et al., (2011)) and a publicly available line (BDSC#24205: *w1118;Mi{ET}IR75dMB04616* – Bloomington *Drosophila* Stock Center, IN-USA). To generate DsuzIR64aand CpomIR64a lines, we used a pIR84a-Gal4 knock-in parental line upon balancing (*w;Bl/CyO;pIR84a-Gal4^KI^* – Grosjean et al., (2011)), which Gal4-knock-in replacing IR84a made an IR84a-knock-out out of it. The final strains tested by SSR had the following genotypes: CpomIR41a1 line: *w;pIR76a-Gal4;pUAS-CpomIR41a1* ; DsuzIR75d line: *w;pUAS-DsuzIR75dHEK/pIR75d-Gal4;IR75d^KO^* ; DsuzIR64a line: *w;pUAS-DsuzIR64a;pIR84a-Gal4* ; CpomIR64a line: *w;pUAS-CpomIR64a;pIR84a-Gal4*. Insects were reared in our facilities at room temperature (19-22 °C) under a 16:8 hours light:dark photoperiod as described in Cattaneo et al., (2022).

### Single sensillum recordings

CpomIR41a1 and DsuzIR75d expressed in the neurons of coeloconic sensilla were tested through single sensillum recordings (SSR). Insects were immobilized as we previously described (Cattaneo et al., 2022). In brief, 3-to-8 day old male flies were inserted in 100 μL pipette tips with only the top half of the head protruding. For each insect, the right antenna of the animal was gently pushed with a glass capillary against a piece of glass. This piece of glass and the pipette tip were fixed with dental wax on a microscope slide. Electrolytically sharpened tungsten electrodes (Harvard Apparatus Ltd, Edenbridge, United Kingdom) were used to penetrate the insect’s body: the reference electrode was manually inserted in the right eye of the fly, while the recording electrode was maneuvered with a DC-3K micromanipulator equipped with a PM-10 piezo translator (Märzhäuser Wetzler GmbH, Wetzler, Germany) and inserted into ac-sensilla. Signals coming from sensory neurons were amplified 10 times with a probe (INR-02, Syntech, Hilversum, the Netherlands), digitally converted through an IDAC-4-USB (Syntech) interface, and visualized and analyzed with the software Autospike v. 3.4 (Syntech). To carry the odorant stimulus, to prevent antennal dryness and to minimize the influence of background odors from the environment, a constant humidified flow of 2.5 L/min charcoal-filtered air was delivered through a glass tube and directed to the preparation. A panel of 23 ligands (Table 1) included control ligands, aimed to indicate choice of the correct sensilla among ac1-4, that were selected based both on findings from Silbering et al., (2011) and upon consulting the DoOR-database of odorant responses (http://neuro.uni-konstanz.de/DoOR/content/DoOR.php – Munch and Galizia (2016), Galizia et al., (2010)). Among possible IR-specific ligands, amines potentially active on IR41a and IR75d were chosen starting from previous deorphanization findings of the *D. melanogaster* orthologues (Herre et al., 2022; Hussain et al., 2016; Silbering et al., 2011). Possible IR64a-activators were chosen by consulting the DoOR – Database of Odorant Responses.

To screen the panel, we adjusted a similar protocol recently adopted in our labs (Cattaneo et al., 2022). In brief, stimuli were diluted either in water or ethanol (Sigma Aldrich, St. Louis, MO-USA) depending on their solubility (Table 1). Stimuli were prepared as done by Silbering et al., (2011), using 20 μL of 1.0% dilutions and 30 μL of 10 μg/μL for phenylacetic acid. As done in our previous experiments (Cattaneo et al., 2022) stimuli aliquots were spread on grade 1 – 20 mm circles filter paper (GE Healthcare Life Science, Little Chalfont, United Kingdom), previously inserted into glass Pasteur pipettes (VWR, Milan, Italy). To minimize possible effects from the solvent, pipettes were let at least 10 minutes after preparation under the fume hood for solvent evaporation. Puffing provided additional 2.5 mL air through the pipette for 0.5 seconds, by inserting the pipette within a side hole of the glass tube directing the humidified air-flow to the antennae. The intensity of the response was quantified by counting all spikes recorded from an individual sensillum as conducted in Silbering et al., (2011) because of the given difficulties in reliably distinguishing spikes from individual neurons (Yao et al., 2005). Spike frequency was calculated by subtracting spikes counted for 0.5 seconds before the stimulus from the number of spikes counted for 0.5 seconds after the stimulus (Δspikes/0.5sec). Responses to compounds of the panel were compared for 5 to 9 replicates, using a single insect as a replicate. To validate significant differences in spike counting, spike frequency of each compound were compared with respective spike frequencies enhanced by the solvent (water/ethanol) by two-tailed paired T-test [::J=0.05] as done in our previous studies (Cattaneo et al., 2022).

To confirm that our recordings were targeting ac4-sensilla, we tested effects on wild type (Oregon) and knock-out insects, both for IR75d (*w;pIR75d-Gal4;IR75d^KO^*and *w;pUAS-DsuzOR75dHEK;IR75d^KO^*) and for IR84a (*w;Bl/CyO;pIR84a-Gal4^KI^*). We then tested effects to the same ligands on transgenic insects expressing DsuzIR75d^HEK^ (*w;pUAS-DsuzIR75d^HEK^/pIR75d-Gal4;IR75d^KO^*), CpomIR41a1 (*w;pIR76a-Gal4;pUAS-CpomIR41a1*) and Dsuz/CpomIR64as (*w;pUAS-DsuzIR64a;pIR84a-Gal4^KI^* and *w;pUAS-CpomIR64a;pIR84a-Gal4^KI^*). Aliquots of ac4-specific ligands enhanced a repeatable and evident neuronal spiking, rather phasic or tonic, depending on the tested ligand, which two-tailed paired T-test comparison with the effect associated with solvents suggested specificity to ac4-sensilla (Supplementary data file 1).

### Dose response SSR experiments

When testing CpomIR41a1 and DsuzIR75d, SSR method was adopted to perform dose-response experiments. For CpomIR41a1 we selected 1,4-diamino butane (putrescine) and N-(3-amminopropil)butan-1,4-diammina (spermidine) among the most active and specific ligands from our SSR-screening (Figure 2 A). On the same insects, a control dose-response experiment was performed by testing hexylamine that we identified to be the most ac4-active ligand for every genotype we tested, most likely reflecting IR76a activation. For DsuzIR75d we selected hexanoic acid, which was the sole ligand active on this subunit from our SSR-screening (Figure 1 B).

To test dose-responses, compounds were diluted in water at [0.1-3.0]%. Depending on the experiment, 1.0-30 μL aliquots were applied on 1-20 mm circles filter paper, in order to provide stimuli, in which neat ligands ranged from about 1.0 nL to about 1.0 μL (doses 0.1 to 90, Supplementary data file 3 – see respective sheet). Spike frequencies were calculated as Δspikes/0.5sec. Normalization for CpomIR41a1 followed similar protocols we have previously adopted (Cattaneo et al., 2022), choosing the effect from saturating doses of putrescine (DOSE 20), which was selected as the compound with the lower molecular weight (MW_putr._ = 88,15 g/mol; MW_sperm._ = 145,25 g/mol) and the highest volatility (Vp_putr._ = 2,33 mmHg; Vp_sperm._ = 0,001 mmHg) against spermidine. Normalization for DsuzIR75d was based on the saturating DOSEs 10 or 20, depending on the experiment. Normalized data were analyzed by Sigma Plot 13.0 (Systat Software Inc., San Jose, CA, USA). Depending on the experiment, responses to selected compounds were compared for 5-13 replicates, considering a replicate as a single insect (Supplementary data file 3).

### Fluorescent in situ hybridization

FISH was performed as we recently reported in Cattaneo et al., (2022). In brief, we generated/synthesized single DIG and FLUO-probes starting from linearized pDONR221-vectors containing DsuzIR64a-coding sequences. A total amount of 1.5 μg of DNA-vector containing DsuzIR64a were linearized with *HpaI* following recommended protocols (New England Biolabs, Ipswich, MA, USA) to be purified in RNAse-free water and checked on agarose gel electrophoresis to verify the linearization of the plasmids. One third of the purified volume (∼ 0.5 μg) was amplified with T7-RNA polymerase (Promega, Madison, WI, USA) integrating DIG-labeled ribonucleotides (BMB Cat. #1277073, Roche, Basel, Switzerland) following recommended protocols (https://www.rockefeller.edu/research/uploads/www.rockefeller.edu/sites/8/2018/10/FISHProtocolKSVRevised.pdf). *D. suzukii* antennae were collected from male and female adult insects of our rearing facility (FORMAS Swedish Research Council-project numbers 2011-390 and 2015-1221). RNA FISH on whole mount antennae were done essentially as described (Saina & Benton, 2013) staining with a single probe for each experiment. Imaging was performed on a Zeiss confocal microscope LSM710 using a 40 × immersion objective; settings were adjusted based on single antennae: DIG-labeled probes staining specific neurons were visualized setting Cy5-laser between 4 and 10% and calibrating gain in a range of 700-900. Staining was compared with male and female FISH-negative control probes for *DsuzIR62a*, which choice as a negative reference was based on non-quantitative RT-PCR results from Walker et al., (2023). Such probes were generated by linearizing pDONR221-vectors containing the DsuzIR62a-coding sequences using *NheI* (New England Biolabs) following protocols described above. The FLUO-labeled RNA-probe that was used to stain *DsuzOrco* as a positive control was the same we adopted in Cattaneo et al., (2022). Neuronal counting was performed using the cell-counter tool of Image J (Fiji, https://imagej.nih.gov/ij/). To identify differences between males and females, neuron numbers were compared with Mann–Whitney U-test (https://www.socscistatistics.com/tests/mannwhitney/) (::J = 0.05, Excel 2016).

### Structural analysis

PDB accessions from DsecIR75b (UniProt A0A1S6EMU3) DmelIR75b (B7Z069), DmelIR64a (Q9VRL4), CpomIR64a (A0A0V0J232), DmelIR84a (Q9VIA5), DmelIR31a (M9PCT4, Isoform C), DmelIR75d (Q9VVU7) and DmelIR8a (Q9W365) were downloaded from Alphafold (alphahold.ebi.cc.uk) and submitted to structural analysis using RasTop (https://www.geneinfinity.org/rastop/). Since the PDB accession of DsuzIR64a is not available, this subunit was not included in the structural analysis. As a preliminary analysis, polypeptide sequence alignment was performed using Muscle (Edgar 2004) and alignment was adjusted with BioEdit (Hall 1999). Based on this, positions of key residues from the LBD were predicted by comparing the subunit of *D. sechellia* IR75d (Prieto-Godino et al., 2017). Polypeptide sequence alignment included orthologues from the IR75cba-clade of *D. melanogaster*, *D.sechellia* and *D. suzukii*, as well as *D. melanogaster* IR64a, IR84a, IR31a, IR75d and IR8a and *C. pomonella* IR64a. Transmembrane domains were predicted with TopCons (Tsirigos et al., 2015), S1/S2 subunits have been assigned according with Prieto-Godino et al., (2017) (Supplementary Figure S2).

### Cloning and heterologous expression in Human Embryonic Kidney cells

The full-length coding sequences of *C. pomonella* IR41a1, IR25a and IR76b and of *D. suzukii* IR8a were obtained from previous studies (Walker et al., 2016; Walker et al., 2023; Ramasamy et al., 2016). Sequences were submitted to GeneArt^TM^ gene synthesis service (Thermos Fischer Scientific, Waltham, MA, USA) requesting integration of the 5’-upstram CACC Kozak-sequence, as done previously (Bobkov et al., 2021; Cattaneo et al., 2017; Crava et al., 2022), codon optimization for expression in human cells (Roberts et al., 2021) and integration into pcDNA3.1 plasmids. To test expression of *D. melanogaster* IR8a+IR84a, the Gibson-assembled combination of these two CDS into pcDNA3.1 was courtesy provided by Dr. Hayden R. Schmidt (International AIDS Vaccine Initiative, USA). Human Embryonic Kidney (HEK293T) cells were grown to semi-confluence in 35-mm Petri dishes containing HEK cell media [Dulbecco’s modified Eagle’s medium containing 10% fetal bovine serum (MP Biomedicals, Solon, OH, United States), 2.0 mM L-glutamine, and 100 mg/ml penicillin/streptomycin (Invitrogen)] at 37°C and 5% CO_2_. For CpomIR41a1-experiments, transient expression was conducted co-transfecting 2.0 μg of pcDNA5/TO-CpomOrco (Cattaneo et al., 2017) with same amounts of pcDNA3.1-CpomIR25a, pcDNA3.1-CpomIR76b and pcDNA3.1-CpomIR41a1. To report expression, 1.0 μg of a separate plasmid DNA carrying the CDS for a blue fluorescent protein (EBFP) was co-transfected [pEBFP2-Nuc, a gift from Robert Campbell, University of Alberta, Alberta, Canada: Ai et al., (2007) -(Addgene plasmid #14893)]. To monitor calcium, 1.0 μg of a separate plasmid DNA carrying the CDS for a genetically-encoded fluorescent Calcium reporter, previously prepared in our labs (pEZT-BH-GCaMP) was co-transfected in substitution of EBFP. To monitor voltage, 1.0 μg of a separate plasmid DNA carrying the CDS for a genetically-encoded voltage indicator ArcLight-Q239 (Jin et al., 2012, AddGene #36856) was co-transfected in substitution of EBFP. Expression of fluorescent reporter genes was under the regulation of the same promoter for Orco/IR genes (CMV). In brief, transfection DNAs was mixed with 3.0 μL FUGENE (Fugent LLC, Middleton, WI-USA) per μg DNA following the recommended protocol to incubate cells overnight for up to 48 h. After incubation, HEK cell media was replaced with 2.0 mL fresh media to incubate cells at 37°C for up to 6-8 additional hours. Part of the cell culture was split in the middle of Mat Tek P35G-1.5-14-C dishes (Ashland, MA USA) as individual cells or small clusters and rinsed at the sides with 2.0 ml fresh HEK media. After splitting, cells were allowed to recover for at least 1 day prior to imaging.

### Imaging Experiments

Activation of HEK293T cells transfected with CpomOrco+IR25a+IR76b+IR41a1 was tested using the same procedures we previously described (Bobkov et al., 2021; Cattaneo et al., 2017; Crava et al., 2022). Mat Tek petri dishes containing HEK293T cells co-transfected with Calcium (GCaMP) or voltage (ArcLightQ239) indicators, underwent to imaging experiments directly upon replacing HEK cell media with 2.0 mL HEK Ca^++^ Ringer (mM: 140 NaCl, 5 KCl, 2 CaCl_2_, 10 HEPES, pH 7.4). To test HEK293T cells for sodium (Na^+^) or potassium (K^+^) permeability, Mat Tek petri dishes were incubated for 1 h at room temperature in 1.0 mL HEK Ca^++^ Ringer, containing either the fluorescent indicator for Na^+^ or for K^+^ (NaTRIUM Green-2 AM, ION Potassium Green-4 AM, ION Indicators, San Marcos, TX USA), prepared at 5-15 mM with 0.06-0.2% Pluronic F-127 (Invitrogen). The buffer was removed after incubation, cells were rinsed with 4 ml fresh HEK Ca^++^ Ringer and placed on the stage of a Zeiss confocal microscope LSM710 using a 20 × objective. Settings were adjusted based on single preparations, visualizing cells using 488-laser at 4%, gain 700-900. Cells were continuously perfused with Ca^++^ Ringer in the course of the experiment using a home-made gravity fed perfusion system.

The perfusion system was constructed by combining two syringes (test VS wash) on a solid tube supporter with connected silicon tubes, and adjusting the height of the syringes to a chronometric flow-rate proximal to 400 μL/min. Silicon tubes terminated in a hand-valve, regulating wash after the stimulus at approximately 65% of the experiment. To provide stimulus on transfected cells, one single silicon tube was directed from this valve to an iron saw terminating with a plastic ClipTip pipette tip (Fisher Scientific), which was placed in the center of the preparation. Extra buffer was gently removed from the Mat Tex petri dish by the use of an additional silicon tube connected to an Ismatek IP-4 peristaltic pump (Fisher Scientific). Fluorescence imaging was performed setting the time-series of the microscope at 100-140 cycles per second. After recording, fluorescent analysis was performed by ImageJ assigning to a sufficient number of green-fluorescent cells a specific region of interest (ROI), for which changes in fluorescence intensity were measured by the tool “ROI manager”. Average fluorescence of the background was subtracted from the average fluorescence of each cell at each single time series. For each cell, basic fluorescence was subtracted from the average fluorescence at each single time series. Changes in fluorescence intensity were expressed as the fractional change in fluorescence intensity (ΔF).

## Acknowledgements

The authors acknowledge the Center for Integrative Genomics, University of Lausanne, for lab and equipment availability. We acknowledge all members of the Benton Lab, University of Lausanne for stimulating discussion, personal communications, material and general availability. We acknowledge Arnaud Paradis from the Cellular Imaging Facility of the University of Lausanne for general advice, availability and general discussion, and the same facility for confocal microscopy availability.

## Author contributions

A.M.C. conceived and designed the experiments, cloned the CDS of IR-subunits, conducted heterologous expression and imaging experiments on HEK-cells, generated transgenic *Drosophila*, performed SSR experiments, *in situ* hybridization experiments and analyzed the data. C.M.C. and W.B.W. implemented the dataset for a preliminary phylogenetic investigation. C.M.C. conceived and designed the preliminary phylogenetic investigation. A.M.C. and W.B.W. conducted structural analysis. A.M.C. supervised the research. A.M.C. and C.M.C. wrote the manuscript. All authors edited and approved the final version of the manuscript.

## Authors approval

All authors have seen and approved the manuscript, and that it hasn’t been accepted or published elsewhere.

## References

Abuin, L., Prieto-Godino, L.L., Pan, H., Gutierrez, C., Huang, L., Jin R. and Benton, R. (2019) In vivo assembly and trafficking of olfactory Ionotropic Receptors. BMC Biology 17:34 10.1186/s12915-019-0651-7

Abuin, L., Bargeton, B., Ulbrich, M. H., Isacoff, E. Y., Kellenberger, S., & Benton, R. (2011). Functional Architecture of Olfactory Ionotropic Glutamate Receptors. Neuron, 69(1), 44–60. 10.1016/j.neuron.2010.11.042

Ai, H. W., Shaner, N. C., Cheng, Z., Tsien, R. Y., & Campbell, R. E. (2007). Exploration of new chromophore structures leads to the identification of improved blue fluorescent proteins. Biochemistry, 46(20), 5904–5910. 10.1021/bi700199g

Ai, M., Min, S., Grosjean, Y., Leblanc, C., Bell, R., Benton, R., & Suh, G. S. (2010). Acid sensing by the Drosophila olfactory system. Nature, 468(7324), 691–695. 10.1038/nature09537

Ai, M. R., Blais, S., Park, J. Y., Min, S., Neubert, T. A., & Suh, G. S. B. (2013). Ionotropic Glutamate Receptors IR64a and IR8a Form a Functional Odorant Receptor Complex *in vivo* in *Drosophila*. Journal of Neuroscience, 33(26), 10741–10749. 10.1523/Jneurosci.5419-12.2013

Armstrong, N., and Gouaux, E. (2000). Mechanisms for activation and antagonism of an AMPA-sensitive glutamate receptor: crystal structures of the GluR2 ligand binding core. Neuron 28, 165–181. 10.1016/S0896-6273(00)00094-5

Benton, R., Vannice, K. S., Gomez-Diaz, C., & Vosshall, L. B. (2009). Variant Ionotropic Glutamate Receptors as Chemosensory Receptors in *Drosophila*. Cell, 136(1), 149–162. 10.1016/j.cell.2008.12.001

Bobkov, Y. V., Walker, W. B., & Cattaneo, A. M. (2021). Altered functional properties of the codling moth Orco mutagenized in the intracellular loop-3. Scientific Reports, 11(1). 10.1038/s41598-021-83024-3

Carey, A., Wang, G., Su, C-Y., Zwiebel, L. & Carlson, J. (2010) Odorant reception in the malaria mosquito *Anopheles gambiae*. Nature 464, 66–71 (2010). 10.1038/nature08834

Cattaneo, A.M. (2018) Current status on the functional characterization of chemosensory receptors of *Cydia pomonella* (Lepidoptera: Tortricidae). Front. Behav. Neurosci. 12:189. 10.3389/fnbeh.2018.00189

Cattaneo, A. M., Gonzalez, F., Bengtsson, J. M., Corey, E. A., Jacquin-Joly, E., Montagne, N., Salvagnin, U., Walker, W. B., Witzgall, P., Anfora, G., & Bobkov, Y. V. (2017). Candidate pheromone receptors of codling moth Cydia pomonella respond to pheromones and kairomones. Scientific Reports, 7. 10.1038/srep41105

Cattaneo, A. M., Witzgall, P., Kwadha, C. A., Becher, P. G., & Walker, W. B. (2022). Heterologous expression and functional characterization of *Drosophila suzukii* OR69a transcript variants unveiled response to kairomones and to a candidate pheromone. Journal of Pest Science. 10.1007/s10340-022-01585-2

Chang, H., Unni, A., Tom, M.T., Llorca, L.C., Brase, S., Bucks, S., Weniger, K., Bisch-Knaden, S., Hansson, B.S., Knaden, M. (2022) Non-redundant odorant detection in a locust. BioRXiv. 10.1101/2022.06.21.496967

Chen, G. Q., Cui, C. H., Mayer, M. L., & Gouaux, E. (1999). Functional characterization of a potassium-selective prokaryotic glutamate receptor. Nature, 402(6763), 817–821. 10.1038/45568

Chiu, J., DeSalle, R., Lam, H. M., Meisel, L., & Coruzzi, G. (1999). Molecular evolution of glutamate receptors: A primitive signaling mechanism that existed before plants and animals diverged. Molecular Biology and Evolution, 16(6), 826–838. 10.1093/oxfordjournals.molbev.a026167

Corcoran, J. A., Sonntag, Y., Andersson, M. N., Johanson, U. & Löfstedt, C. (2018) Endogenous insensitivity to the Orco agonist VUAA1 reveals novel olfactory receptor complex properties in the specialist fly *Mayetiola destructor*. Sci. Rep. 8, 3489. 10.1038/s41598-018-21631-3

Crava, C. M., Bobkov, Y. V., Sollai, G., Anfora, G., Crnjar, R., & Cattaneo, A. M. (2022). Chemosensory Receptors in the Larval Maxilla of *Papilio hospiton*. Frontiers in Ecology and Evolution, 9. 10.3389/fevo.2021.795994

Croset, V., Rytz, R., Cummins, S. F., Budd, A., Brawand, D., Kaessmann, H., Gibson, T. J., & Benton, R. (2010). Ancient Protostome Origin of Chemosensory Ionotropic Glutamate Receptors and the Evolution of Insect Taste and Olfaction. Plos Genetics, 6(8). 10.1371/journal.pgen.1001064

De Fouchier, A., Walker III, W. B., Montagné, N., Steiner, C., Binyameen, M., Schlyter, F., … & Jacquin-Joly, E. (2017). Functional evolution of Lepidoptera olfactory receptors revealed by deorphanization of a moth repertoire. Nat comm, 8(1), 15709. 10.1038/ncomms15709

Edgar, R. C. (2004). MUSCLE: multiple sequence alignment with high accuracy and high throughput. Nucleic Acids Research, 32(5), 1792–1797. 10.1093/nar/gkh340

Egger, B., van Giesen, L., Moraru, M., & Sprecher, S. G. (2013). *In vitro* imaging of primary neural cell culture from *Drosophila*. Nat Protoc, 8(5), 958–965. 10.1038/nprot.2013.052

Enjin, A., Zaharieva, E. E., Frank, D. D., Mansourian, S., Suh, G. S., Gallio, M., & Stensmyr, M. C. (2016). Humidity Sensing in *Drosophila*. Current Biology, 26(10), 1352–1358. 10.1016/j.cub.2016.03.049

Eyun, S.I., Soh, H.Y., Posavi, M., Munro, J.B., Hughes, D.S., Murali, S.C., Qu, J., Dugan, S., Lee, S.L., Chao, H. and Dinh, H., 2017. Evolutionary history of chemosensory-related gene families across the Arthropoda. Molecular biology and evolution, 34(8), pp.1838–1862. 10.1093/molbev/msx147

Galizia, C. G., Munch, D., Strauch, M., Nissler, A., & Ma, S. (2010). Integrating heterogeneous odor response data into a common response model: A DoOR to the complete olfactome. Chemical Senses, 35(7), 551–563. 10.1093/chemse/bjq042

Gereau, R. W., & Swanson, G. T. (2008). The Glutamate Receptors. https://link.springer.com/book/10.1007/978-1-59745-055-3

German PF, van der Poel S, Carraher C, Kralicek AV, Newcomb RD. 2013. Insights into subunit interactions within the insect olfactory receptor complex using FRET. Insect Biochem Mol Biol. 43:138–145. 10.1016/j.ibmb.2012.11.002

Gonzalez, F., Witzgall, P., & Walker, W. B. (2016). Protocol for Heterologous Expression of Insect Odourant Receptors in *Drosophila*. Frontiers in Ecology and Evolution, 4. 10.3389/fevo.2016.00024

Gorter, J., Jagadeesh, S., Gahr, C., Boonekamp, J.J., Levine, J.D. & Billeter, J-C. (2016) The nutritional and hedonic value of food modulate sexual receptivity in *Drosophila melanogaster* females. Sci Rep 6, 19441 (2016). 10.1038/srep19441

Grosjean, Y., Rytz, R., Farine, J. P., Abuin, L., Cortot, J., Jefferis, G. S., & Benton, R. (2011). An olfactory receptor for food-derived odours promotes male courtship in *Drosophila*. Nature, 478(7368), 236–240. 10.1038/nature10428

Grosse-Wilde, E., Gohl, T., Bouche, E., Breer, H. and Krieger, J. 2007. Candidate pheromone receptors provide the basis for the response of distinct antennal neurons to pheromonal compounds. European Journal of Neuroscience, 25, 2364–2373. 10.1111/j.1460-9568.2007.05512.x

Grosse-Wilde, E., Svatos, A. & Krieger, J. 2006. A pheromone-binding protein mediates the bombykol-induced activation of a pheromone receptor *in vitro*. Chemical Senses, 31, 547–555. 10.1093/chemse/bjj059

Guo, M., Du, L., Chen, Q., Feng, Y., Zhang, J., Zhang, X., Tian, K., Cao, S., Huang, T., Jacquin-Joly, E., Wang, G & Liu, Y. (2021). Odorant receptors for detecting flowering plant cues are functionally conserved across moths and butterflies. Molecular biology and evolution, 38(4), 1413–1427. 10.1093/molbev/msaa300

Gupta, A., Singh, S.S., Mittal, A.M., Singh, P., Goyal, S., Kannan, K.R., Gupta, A.K., Gupta N. (2022). Mosquito Olfactory Response Ensemble enables pattern discovery by curating a behavioral and electrophysiological response database iScience 25(3), 103938. 10.1016/j.isci.2022.103938

Hall, T. A. (1999). BioEdit: a user-friendly biological sequence alignment editor and analysis program for Windows 95/98/NT. Nucleic Acids Symposium Series, 41, 95–98.

Herre, M., Goldman, O. V., Lu, T. C., Caballero-Vidal, G., Qi, Y. Y., Gilbert, Z. N., Gong, Z. Y., Morita, T., Rahiel, S., Ghaninia, M., Ignell, R., Matthews, B. J., Li, H. J., Vosshall, L. B., & Younger, M. A. (2022). Non-canonical odor coding in the mosquito. Cell, 185(17), 3104-+. 10.1016/j.cell.2022.07.024

Hopf, T.A., Morinaga, S., Ihara, S., Touhara, K., Marks, D.S. and Benton R. (2015) Amino acid coevolution reveals three-dimensional structure and functional domains of insect odorant receptors. Nat. Comm. Volume 6: 6077. 10.1038/ncomms7077

Hou, X., Zhang, D. D., Yuvaraj, J.K., Corcoran, J.A., Andersson, M. N., & Lofstedt, C. (2020). Functional characterization of odorant receptors from the moth *Eriocrania semipurpurella*: A comparison of results in the *Xenopus* oocyte and HEK cell systems. Ins. Biochem. and Mol. Biol. 117, 103289. 10.1016/j.ibmb.2019.103289

Hou, X. Q., Zhang, D. D., Powell, D., Wang, H. L., Andersson, M. N., & Lofstedt, C. (2022). Ionotropic receptors in the turnip moth *Agrotis segetum* respond to repellent medium-chain fatty acids. Bmc Biology, 20(1), 34. 10.1186/s12915-022-01235-0

Hueston, C. E., Olsen, D., Li, Q., Okuwa, S., Peng, B., Wu, J., & Volkan, P. C. (2016). Chromatin modulatory proteins and olfactory receptor signaling in the refinement and maintenance of fruitless expression in olfactory receptor neurons. PLoS biology, 14(4), e1002443. 10.1371/journal.pbio.1002443

Hussain, A., Zhang, M., Ucpunar, H. K., Svensson, T., Quillery, E., Gompel, N., Ignell, R., & Kadow, I. C. G. (2016). Ionotropic Chemosensory Receptors Mediate the Taste and Smell of Polyamines. Plos Biology, 14(5). 10.1371/journal.pbio.1002454

Jin, L., Han, Z., Platisa, J., Wooltorton, J. R., Cohen, L. B., & Pieribone, V. A. (2012). Single action potentials and subthreshold electrical events imaged in neurons with a fluorescent protein voltage probe. Neuron, 75(5), 779–785. 10.1016/j.neuron.2012.06.040

Koh, T. W., He, Z., Gorur-Shandilya, S., Menuz, K., Larter, N. K., Stewart, S., & Carlson, J. R. (2014). The *Drosophila* IR20a Clade of Ionotropic Receptors Are Candidate Taste and Pheromone Receptors. Neuron, 83(4), 850–865. 10.1016/j.neuron.2014.07.012

Kurtovic, A., Widmer, A., & Dickson, B. J. (2007). A single class of olfactory neurons mediates behavioural responses to a *Drosophila* sex pheromone. Nature, 446(7135), 542–546. 10.1038/nature05672

Lam, H. M., Chiu, J., Hsieh, M. H., Meisel, L., Oliveira, I. C., Shin, M., & Coruzzi, G. (1998). Glutamate-receptor genes in plants. Nature, 396(6707), 125–126. 10.1038/24066

Llopis-Giménez, A., Caballero-Vidal, G., Jacquin-Joly, E., Crava, C.M., Herrero, S. (2021) Baculovirus infection affects caterpillar chemoperception. Ins Biochem and Mol Physiol 138: 103648 10.1016/j.ibmb.2021.103648

Miazzi, F., Hoyer, C., Sachse, S., Knaden, M, Wicher, D., Hansson, B.S., Lavista-Llanos, S. (2019) Optimization of insect odorant receptor trafficking and functional expression via transient transfection in HEK293 cells. Chem Senses 44(9):673–682. 10.1093/chemse/bjz062

Min, S., Ai, M., Shin, S. A., & Suh, G. S. (2013). Dedicated olfactory neurons mediating attraction behavior to ammonia and amines in *Drosophila*. Proc Natl Acad Sci U S A, 110(14), E1321–1329. 10.1073/pnas.1215680110

Munch, D., & Galizia, C. G. (2016). DoOR 2.0--Comprehensive Mapping of *Drosophila* melanogaster Odorant Responses. Sci Rep, 6, 21841. 10.1038/srep21841

Ni, L. (2021) The Structure and Function of Ionotropic Receptors in *Drosophila*. Front. Mol. Neurosci. Sec. Neuroplasticity and Development. 13. 10.3389/fnmol.2020.638839

Pitts, R. J., Derryberry, S. L., Zhang, Z., & Zwiebel, L. J. (2017). Variant Ionotropic Receptors in the Malaria Vector Mosquito *Anopheles gambiae* Tuned to Amines and Carboxylic Acids. Sci Rep, 7, 40297. 10.1038/srep40297

Prieto-Godino, L.L., Schmidt, H.R., Benton R. (2021) Molecular reconstruction of recurrent evolutionary switching in olfactory receptor specificity. Elife 10:e69732. 10.7554/eLife.69732

Prieto-Godino, L. L., Rytz, R., Cruchet, S., Bargeton, B., Abuin, L., Silbering, A. F., Ruta, V., Dal Peraro, M., & Benton, R. (2017). Evolution of Acid-Sensing Olfactory Circuits in Drosophilids. Neuron, 93(3), 661–676 e666. 10.1016/j.neuron.2016.12.024

Prieto-Godino, L.L, Rytz, R., Bargeton, B., Abuin, l., Arguello, J.R., Dal Peraro, M. and Benton, R. (2016) Olfactory receptor pseudo-pseudogenes. Nature 539, 93–97. 10.1038/nature19824

Ramasamy, S., Ometto, L., Crava, C. M., Revadi, S., Kaur, R., Horner, D. S., Pisani, D., Dekker, T., Anfora, G., & Rota-Stabelli, O. (2016). The Evolution of Olfactory Gene Families in *Drosophila* and the Genomic Basis of chemical-Ecological Adaptation in *Drosophila suzukii*. Genome Biology and Evolution, 8(8), 2297–2311. 10.1093/gbe/evw160

Ray, G., Huff, R. M., Castillo, J. S., Bellantuono, A. J., DeGennaro, M., & Pitts, R. J. (2023). Carboxylic acids that drive mosquito attraction to humans activate ionotropic receptors. PLOS Neglected Tropical Diseases, 17(6), e0011402. 10.1371/journal.pntd.0011402

Roberts, R. E., Yuvaraj, J. K., and Andersson, M. N. (2021). Codon Optimization of Insect Odorant Receptor Genes May Increase Their Stable Expression for Functional Characterization in HEK293 Cells. Front. Cell. Neurosci., 15. 10.3389/fncel.2021.744401

Saina, M., & Benton, R. (2013). Visualizing Olfactory Receptor Expression and Localization in Drosophila. In C. J. Crasto (Ed.), Olfactory Receptors Methods and Protocols. 10.1007/978-1-62703-377-0

Shan, S., Wang, S-N., Song, X., Khashaveh, A., Lu, Z-Y., Hussain Dhiloo, K., Li, R-J., Gao, X-W., Zhang, Y-J. (2019) Antennal ionotropic receptors IR64a1 and IR64a2 of the parasitoid wasp *Microplitis mediator* (Hymenoptera: Braconidate) collaboratively perceive habitat and host cues. Insect Biochemistry and Molecular Biology Volume 114:103204 10.1016/j.ibmb.2019.103204

Silbering, A. F., Rytz, R., Grosjean, Y., Abuin, L., Ramdya, P., Jefferis, G. S. X. E., & Benton, R. (2011). Complementary Function and Integrated Wiring of the Evolutionarily Distinct *Drosophila* Olfactory Subsystems. Journal of Neuroscience, 31(38), 13357–13375. 10.1523/Jneurosci.2360-11.2011

Soffan, A., Subandiyah, S., Makino, H., Watanabe, T., & Horiike, T. (2018). Evolutionary Analysis of the Highly Conserved Insect Odorant Coreceptor (Orco) Revealed a Positive Selection Mode, Implying Functional Flexibility. J Insect Sci, 18(6). 10.1093/jisesa/iey120

Stewart, S., Koh, T. W., Ghosh, A. C., & Carlson, J. R. (2015). Candidate ionotropic taste receptors in the *Drosophila* larva. Proceedings of the National Academy of Sciences of the United States of America, 112(14), 4195–4201. 10.1073/pnas.1503292112

Swanson, G. T., Kamboj, S. K., & Cull-Candy, S. G. (1997). Single-channel properties of recombinant AMPA receptors depend on RNA editing, splice variation, and subunit composition. Journal of Neuroscience, 17(1), 58–69. 10.1523/JNEUROSCI.17-01-00058.1997

Syed, Z., Kopp, A., Kimbrell, D. A., & Leal, W. S. (2010). Bombykol receptors in the silkworm moth and the fruit fly. Proceedings of the National Academy of Sciences, 107(20), 9436–9439. 10.1073/pnas.1003881107

Syed, Z., Ishida, Y., Taylor, K., Kimbrell, D. A., & Leal, W. S. (2006). Pheromone reception in fruit flies expressing a moth’s odorant receptor. Proc Natl Acad Sci U S A, 103(44), 16538–16543. 10.1073/pnas.0607874103

Tauber, J. M., Brown, E. B., Li, Y. Y., Yurgel, M. E., Masek, P., & Keene, A. C. (2017). A subset of sweet-sensing neurons identified by IR56d are necessary and sufficient for fatty acid taste. Plos Genetics, 13(11). 10.1371/journal.pgen.1007059

Tsirigos, K. D., Peters, C., Shu, N., Kall, L., & Elofsson, A. (2015). The TOPCONS web server for consensus prediction of membrane protein topology and signal peptides. Nucleic Acids Research, 43(W1), W401–W407. 10.1093/nar/gkv485

Turner, R. M., Derryberry, S.L., Kumar, B.N., Brittain, T., Zwiebel, L.J., Newcomb, R.D., Christie, D.L. (2014) Mutational analysis of cysteine residues of the insect odorant co-receptor (Orco) from *Drosophila melanogaster* reveals differential effects on agonist-and odorant-tuning receptor-dependent activation. J. Biol. Chem. 289(46), 31837–31845. 10.1074/jbc.M114.603993

Ui, K., Nishihara, S., Sakuma, M., Togashi, S., Ueda, R., Miyata, Y., and Miyake, T. (1994) Newly established cell lines from *Drosophila* larval CNS express neural specific characteristics. In Vitro Cell Dev Biol – Animal 30, 209–216 (1994). 10.1007/BF02632042

van Giesen, L., & Garrity, P. A. (2017). More than meets the IR: the expanding roles of variant Ionotropic Glutamate Receptors in sensing odor, taste, temperature and moisture. Version 1. F1000Res., 6(1753). 10.12688/f1000research.12013.1

Vulpe, A. and Menuz, K. (2021) Ir76b is a Co-receptor for Amine Responses in *Drosophila* Olfactory Neurons. Front Cell Neurosci . 15:759238. 10.3389/fncel.2021.759238

Walker, W. B., 3rd, Gonzalez, F., Garczynski, S. F., & Witzgall, P. (2016). The chemosensory receptors of codling moth Cydia pomonella-expression in larvae and adults. Sci Rep, 6, 23518. 10.1038/srep23518

Walker, W. B., 3rd, Mori, B. A., Cattaneo, A. M., Gonzalez, F., Witzgall, P., & Becher, P. G. (2023). Comparative transcriptomic assessment of the chemosensory receptor repertoire of Drosophila suzukii adult and larval olfactory organs. Comp Biochem Physiol Part D Genomics Proteomics, 45, 101049. 10.1016/j.cbd.2022.101049

Wang, B., Liu, Y., He, K. and Wang, G. (2016) Comparison of research methods for functional characterization of insect olfactory receptors. Sci Rep 6, 32806 (2016). 10.1038/srep32806

Wicher, D., & Miazzi, F. (2021). Functional properties of insect olfactory receptors: ionotropic receptors and odorant receptors. Cell and Tissue Research, 383(1), 7–19. 10.1007/s00441-020-03363-x

Xu, P., Choo, Y. M., De La Rosa, A., & Leal, W. S. (2014). Mosquito odorant receptor for DEET and methyl jasmonate. Proceedings of the National Academy of Sciences, 111(46), 16592–16597. 10.1073/pnas.141724411

Yao, C. A., Ignell, R., & Carlson, J. R. (2005). Chemosensory coding by neurons in the coeloconic sensilla of the *Drosophila* antenna. Journal of Neuroscience, 25(37), 8359–8367. 10.1523/JNEUROSCI.2432-05.2005.

Yin, N-N., Nuo, S-M., Xiao, H-Y., Zhao, Y-J., Zhu, J-Y., Liu, N-Y. (2021) The ionotropic receptor gene family in Lepidoptera and Trichoptera: Annotation, evolutionary and functional perspectives. Genomics, 113(1) Part 2, 601-612. 10.1016/j.ygeno.2020.09.056

You, Y., Smith, D.P., Lv, M., Zhang, L. (2016) A broadly tuned odorant receptor in neurons of trichoid sensilla in locust, *Locusta migratoria*. Insect Biochem Mol Biol 79:66–72. 10.1016/j.ibmb.2016.10.008

