## Supplementary figures and images for "Heterologous expression of insect IRs in transgenic *Drosophila melanogaster*"

### Supplementary Figure S1

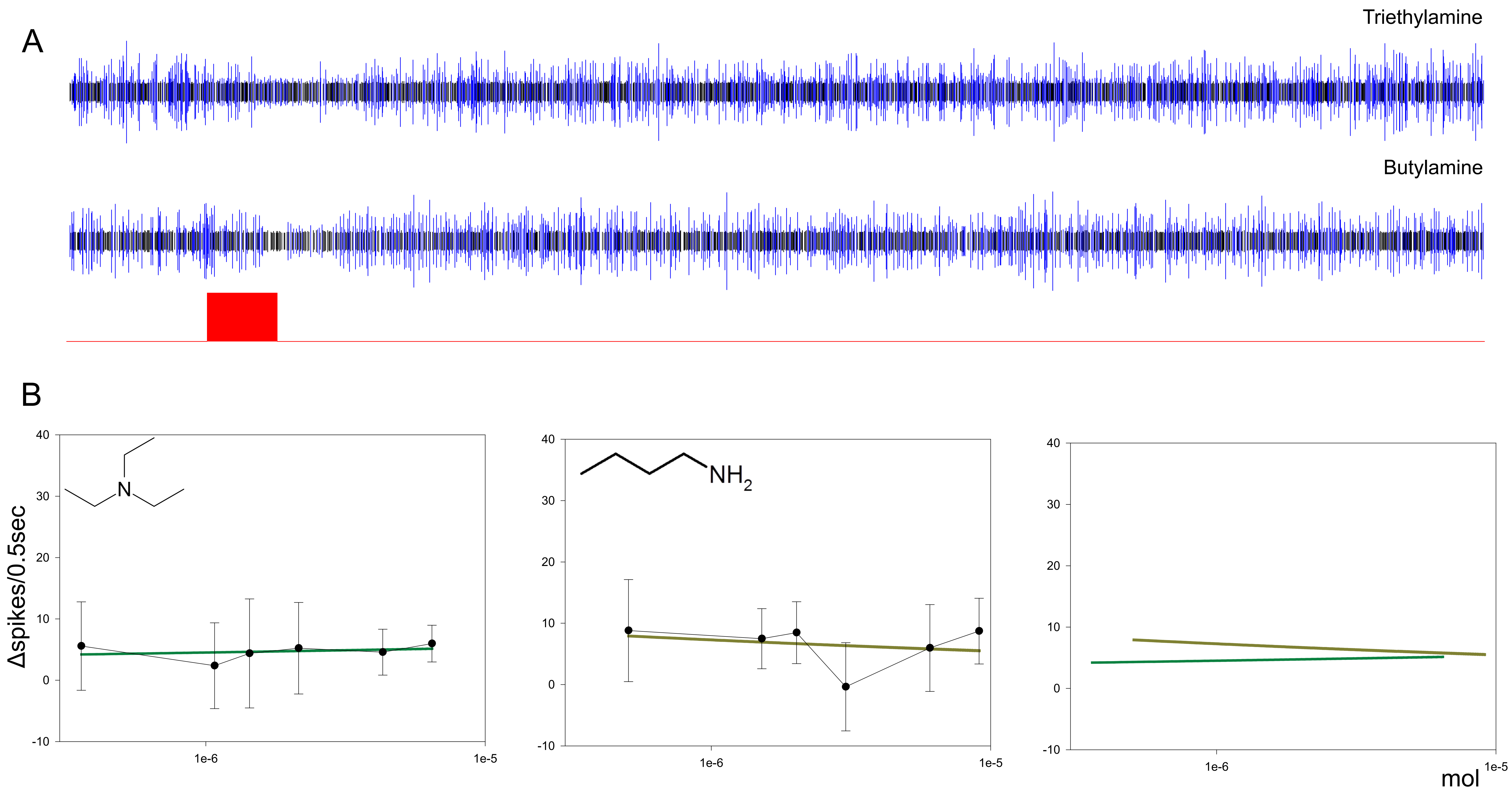

### Supplementary Figure S2

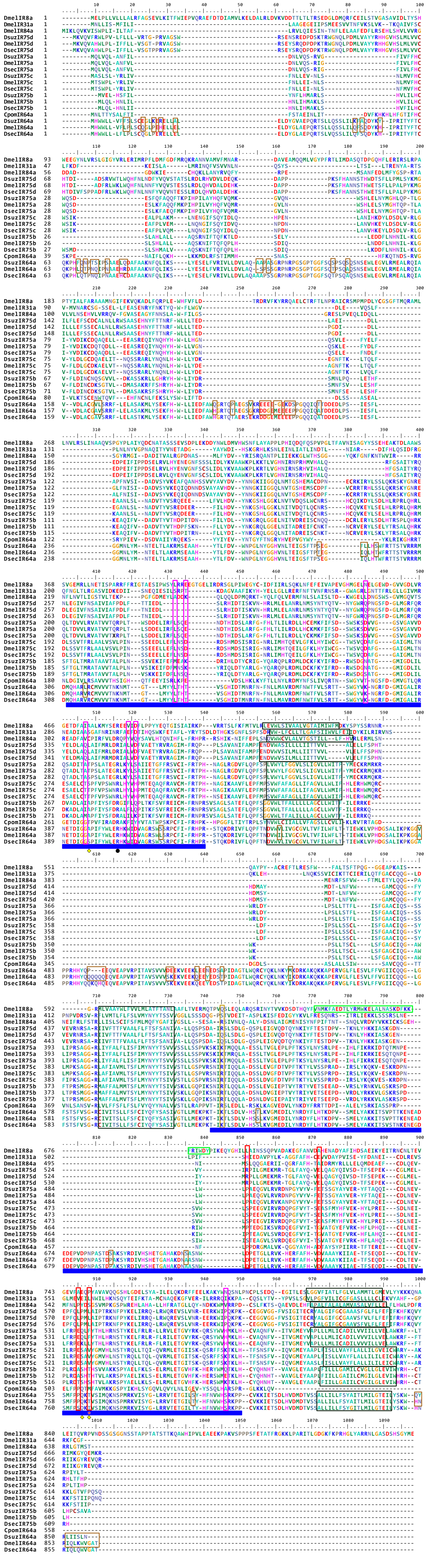
