## Supplementary Figure S3 for "Heterologous expression of insect IRs in transgenic *Drosophila melanogaster*"

### CpomIR41a lines

*w ; pIR76a-Gal4 ;* $\frac{TM2}{TM6B}$ X *w ;* $\frac{Bl}{CyO}$ *; pUAS-**CpomIR41a1*** → *w ; pIR76a-Gal4 ; pUAS-**CpomIR41a1***

### DsuzIR75d$^{HEK}$ lines

*w ; pIR75d-Gal4 ;* $\frac{TM2}{TM6B}$ X *w ;* $\frac{Bl}{CyO}$ *;* $IR75d^{KO}$ → *w ; pIR75d-Gal4 ;* $IR75d^{KO}$

X → *w ;* $\frac{pIR75d\text{-}Gal4}{pUAS\text{-}\mathbf{DsuzIR75d}^{HEK}}$ *;* $IR75d^{KO}$

*w ; pUAS-**DsuzIR75d***$^{HEK}$ *;* $\frac{TM2}{TM6B}$ X *w ;* $\frac{Bl}{CyO}$ *;* $IR75d^{KO}$ → *w ; pUAS-**DsuzIR75d***$^{HEK}$ *;* $IR75d^{KO}$

### Dsuz/CpomIR64a lines

*w ; pUAS-**(Dsuz/Cpom)IR64a** ;* $\frac{TM2}{TM6B}$ X *w ;* $\frac{Bl}{CyO}$ *;* $pIR84aGal4^{KI}$ → *w ; pUAS-**(Dsuz/Cpom)IR64a** ;* $pIR84aGal4^{KI}$
